# Multiple clustered centrosomes in antigen-presenting cells foster T cell activation without MTOC polarization

**DOI:** 10.1101/2024.07.18.604057

**Authors:** Isabel Stötzel, Ann-Kathrin Weier, Apurba Sarkar, Subhendu Som, Peter Konopka, Eliška Miková, Jan Böthling, Mirka Homrich, Laura Schaedel, Uli Kazmaier, Konstantinos Symeonidis, Zeinab Abdullah, Stefan Uderhardt, Miroslav Hons, Raja Paul, Heiko Rieger, Eva Kiermaier

## Abstract

Cellular polarization plays a pivotal role in regulating immunological processes and is often associated with centrosome reorientation. During immune synapse (IS) formation centrosome repositioning in lymphocytes assists in T cell activation. While a single centrosome, consisting of two centrioles, is present in T cells, antigen-presenting cells (APCs) such as dendritic cells (DCs) amplify centrioles during maturation leading to increased centrosome numbers upon immune activation. How centrosome amplification in DCs affects IS formation and T cell activation is unclear. In this study, we combine experimental data with mathematical and computational modelling to provide evidence that centrosome amplification in DCs enhances antigen-specific T cell activation. Extra centrioles in DCs form active centrosomes, which cluster during DC-T cell interactions and unlike in T cells, localize close to the cell center. Perturbing either centriole numbers or centrosome configuration in DCs results in impaired T cell activation. Collectively, our results highlight a crucial role for centrosome amplification and optimal centrosome positioning in APCs for controlling T cell responses.

## Introduction

Centrosome repositioning adjacent to the cell cortex is a prerequisite for various fundamental cellular processes such as polarized secretion, directional migration, asymmetric cell division and immune responses (Grill & Hyman, 2005; Li & Gundersen, 2008; Luxton & Gundersen, 2011; Martín-Cófreces *et al*, 2014). T cells are critical cellular players of the adaptive immune system, which combat specific pathogens and damaged body cells. A defining event during T cell activation constitutes the establishment of the IS at the T cell-APC contact site. The formation of the IS starts with the recognition of a cognate antigen by the T cell receptor (TCR), which is presented via major histocompatibility (MHC) complexes on the surface of the APC. Concomitantly, T cell receptors, integrins and co-stimulatory receptors engage with each other at the cell-cell contact site to form a series of supramolecular activation clusters (SMAC), which segregate into radial symmetric zones facing the APC (Grakoui *et al*, 1999; Shaw & Dustin, 1997). Structurally, T cell receptors and associated kinases cluster in the central area (referred to as the central SMAC), while adhesion receptors, actin as well as actin-interacting proteins, rearrange in surrounding rings referred to as the peripheral and distal SMAC (Monks *et al*, 1998).

Formation of the IS is further coupled to major reorganization of the T cells’ microtubule (MT) cytoskeleton: antigen recognition results in the repositioning of the T cell’s centrosome from the uropod to a position adjacent to the IS (Kupfer & Dennert, 1984; Kupfer *et al*, 1987). Centrosome reorientation toward the IS strictly depends on antigen recognition by the cognate T cell receptor (Kupfer & Singer, 1989) and is accompanied by the movement of other organelles such as the Golgi apparatus toward the IS (Kupfer *et al*, 1986, 1985). Polarization of Golgi-derived vesicles is considered to facilitate directional vesicle release toward the target cell (Kupfer *et al*, 1985; Stinchcombe *et al*, 2006). Moreover, MTOC reorientation in T cells has been demonstrated to be required for sustained T cell receptor signaling downstream of IS formation (Martín-Cófreces *et al*, 2008).

Most studies of the IS have focused on those formed between T and B cells. Yet, B cells have a rather week T cell priming capacity (Gunzer *et al*, 2004). DCs represent the most potent APCs of the innate immune system, which have the unique ability to activate T cells *in vivo* (Steinman & Cohn, 1973). They are strategically located in peripheral tissues, where they patrol the environment for invading microbial pathogens. Upon antigen encounter DCs become highly migratory and translocate to draining lymph nodes in order to present peripherally acquired and processed antigens to naïve T cells. Besides antigen presentation, DCs shape T cell responses by secreting large amounts of soluble cytokines that activate T cells (Banchereau *et al*, 2000).

While the tubulin-based cytoskeleton has been widely studied in T cells during IS formation and T cell receptor signaling, little is known about the APC side of the IS. Centrosome polarization has been reported in DCs forming antigen-specific contacts with CD8^+^ T cells in order to allow targeted delivery of T cell stimulatory molecules to the IS (Pulecio *et al*, 2010). In this context, MTs are implicated in playing a crucial role in transporting cargo from the endoplasmatic reticulum (ER) and the Golgi complex to the cell surface (reviewed by Fourriere *et al*, 2020). Moreover, MTs are involved in exocytosis, the final step of secretory protein transport, in which they assist in establishing a localized fusion machinery thereby restricting exocytosis to particular plasma mebrane domains (Fourriere *et al*, 2019).

In leukocytes, MT nucleation primarily takes place at the centrosome (Kopf & Kiermaier, 2021), which consists of two barrel-shaped centrioles that are connected through a flexible linker and surrounded by a proteinaceous matrix referred to as pericentriolar material (PCM) (Bornens, 2012; Paintrand *et al*, 1992). Typically, centrosome numbers are tightly regulated during the cell cycle leading to one centrosome (two centrioles) in G1 phase and two centrosomes (4 centrioles) prior to mitosis (Fırat-Karalar & Stearns, 2014; Nigg & Holland, 2018; Pereira *et al*, 2021). Of note, centrosome numbers and protein composition can vary during organismal development and differentiation into specialized cell types leading to cells that contain either no centrosome or multiple (Muroyama & Lechler, 2017; Meyer-Gerards & Bazzi, 2024; Carden *et al*, 2023). During infection, antigen encounter provokes DCs to enter a robust cell cycle arrest which can be accompanied by the acquisition of extra centrosomes in G1 phase (Weier *et al*, 2022). Extra centrosomes in DCs nucleate MT filaments and promote persistent migration of cells toward chemotactic cues. Moreover, DCs with multiple centrosomes exhibit a higher capacity for secreting T cell stimulatory molecules and priming of CD4^+^ T cells. Yet, the underlying molecular mechanism(s) of centrosome-mediated enhancement of T cell activation remain unsettled. In particular, we lack information about the organization of multi-numerous centrosomes in APCs during the formation of antigen-specific synapses and how centrosome configuration in APCs affects T cell activation.

In this study, we investigate centrosome organization in murine DCs upon formation of antigen-specific T helper synapses in fixed and living samples. We employ an interdisciplinary approach combining experimental data with computational modeling to study optimal centrosome positioning in APCs and how centrosome numbers, integrity and orientation impact efficient T cell activation.

## Results

### Centrosome and MT integrity in DCs are required for efficient T cell activation

While centrosome function is well established in T cells, its role in APCs is poorly understood. To address whether an intact centrosome in APCs is a prerequisite for efficient T cell activation, we first sought to pharmacologically interfere with centrosome integrity specifically in APCs prior to T cell conjunction. To this end, we used antigen-presenting DCs as a model cell type for studying antigen-specific T cell activation. To obtain sufficiently large numbers of cells, we generated DCs from the bone marrow of Centrin-2 (CETN2)-GFP expressing mice in the presence of granulocyte-macrophage colony-stimulating factor (GM-CSF) and stimulated the cells with lipopolysaccharide (LPS) overnight to induce cell maturation. As mature bone marrow-derived DCs (BMDCs) are a heterogenous population of cells consisting of diploid and tetraploid cells as a result of an incomplete mitosis (Weier *et al*, 2022), we separated BMDCs based on DNA content into diploid (2N) and tetraploid (4N) cells and concentrated exclusively on the 2N cell fraction for all further experiments (**Fig. 1A**). Sorted 2N DCs were further loaded or not loaded with different concentrations of the model antigen ovalbumin-peptide (OVAp) and incubated with naive CD4^+^, OVA-specific T cells, which express a transgenic T cell receptor recognizing the OVAp (OT-II T cells). Under these experimental conditions, OVAp-loaded DCs efficiently activated naive T cells as measured by surface levels of early activation markers such as CD69 and CD62L downregulation as well as T cell proliferation assessed by proliferation-mediated dilution of the fluorescent dye carboxyfluorescein succinimidyl ester (CFSE) (Quah *et al*, 2007) (**Fig. EV1A-D**).

**Figure 1.**
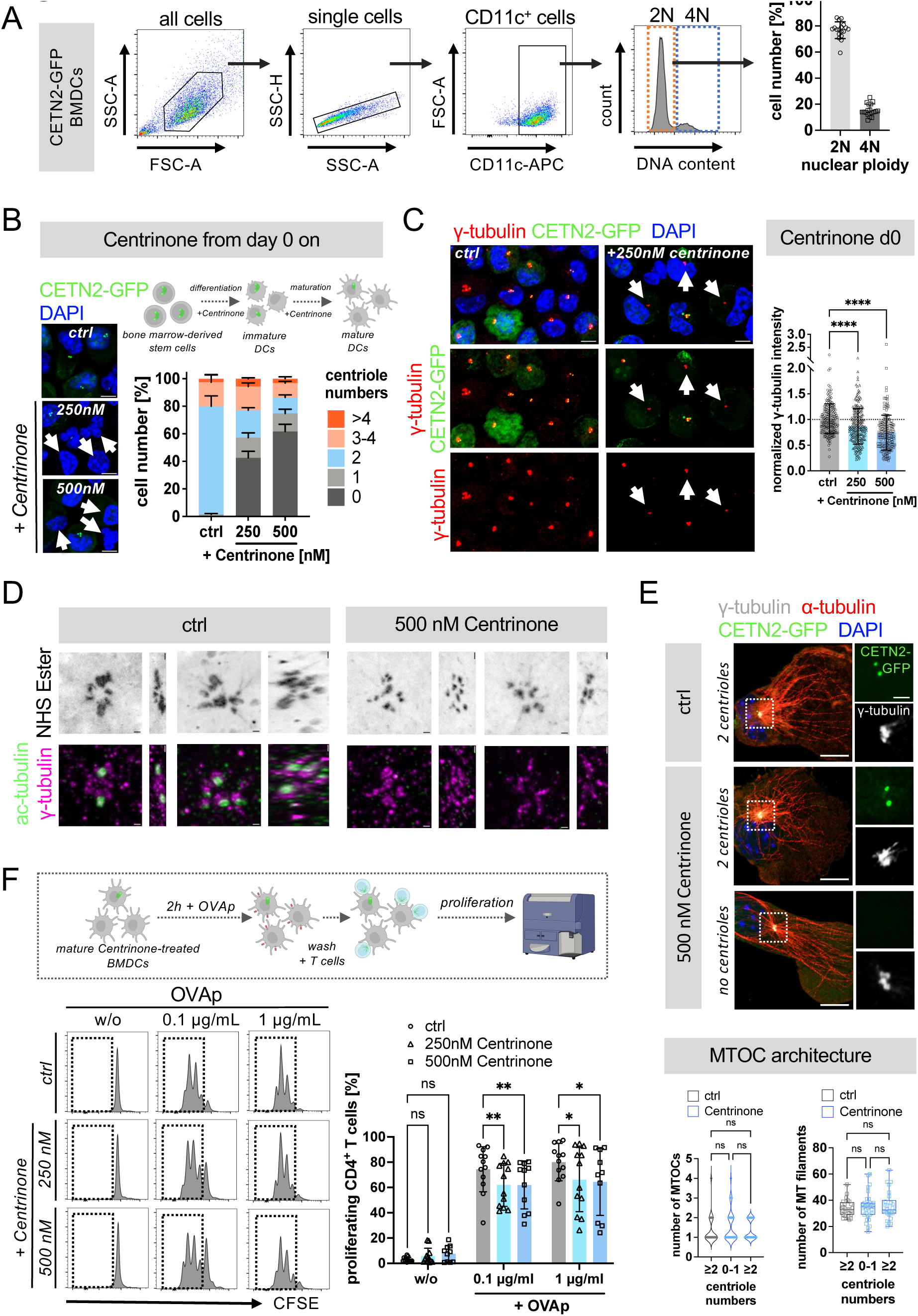
An intact centrosome in DCs is required for efficient T cell activation. (**A**) DNA staining of mature CETN2-GFP expressing BMDCs to determine nuclear ploidy. Left: gating strategy for identification of 2N and 4N DCs and histogram of DNA content distribution of CD11c^+^ cells. Right: quantification of 2N and 4N cells according to DNA content. Graph displays mean values ± s.d. of 18 independent experiments. (**B**) Inhibition of PLK4 activity by Centrinone. CETN2-GFP expressing BMDCs were treated with either 250 or 500 nM Centrinone during differentiation (from day 0 on) and centriole numbers were determined according to CETN2-GFP^+^ foci in mature BMDCs. Left: confocal images of Centrinone-treated and control cells. Merged channels of CETN2-GFP (green) and DAPI (blue) are shown. White arrowheads indicate cells without centrioles. Scale bars, 5 *μ*m. Right: quantification of centriole numbers after Centrinone treatment. Graph displays mean values ± s.d. of three independent experiments with at least *N* = 200 cells analyzed per condition. Picture created with BioRender. (**C**) Immunostaining of mature CETN2-GFP BMDCs against γ-tubulin after 250 nM Centrinone treatment. Merged and individual channels of CETN2-GFP (green), γ-tubulin (red) and DAPI (blue) are shown. Scale bars, 5 *μ*m. White arrowheads indicate cells without centrioles but with prominent γ-tubulin foci. Right: quantification of γ-tubulin signal intensity in mature CETN2-GFP expressing BMDCs after Centrinone treatment. Graph shows normalized values relative to cells with two centrioles ± s.d. Each data point represents one cell derived from one representative experiment out of three independent experiments. Dotted line drawn at 1.0. ****, *P* < 0.0001 (one-way Anova with Dunnett’s multiple comparisons). (**D**) Expansion microscopy of mature control or Centrinone-treated (500nM) CETN2-GFP BMDCs. Top panels: NHS-Ester staining. Bottom panels: merged channels of acetylated (ac)-tubulin (green) and γ-tubulin (magenta). Images are shown as top view and right view. Scale bars, 1 µm. (**E**) Immunostaining of MTs in Centrinone-treated and control cells. Upper panel: mature CETN2-GFP (green) BMDCs were confined under agarose and immunostained against γ-tubulin (white) and α-tubulin (red). Nuclei were counterstained with DAPI (blue). Scale bars, 10 µm. Magnifications of the indicated regions show individual channels of CETN2-GFP (green) and γ-tubulin (white). Scale bars, 2 µm. Lower panel: quantification of MTOCs (left) and MT filaments emanating from defined regions around centrosomes (right) in mature CETN2-GFP expressing BMDCs after Centrinone treatment. Left graph shows median and distribution of data points of at least three independent experiments. ns, non-significant (Kruskal-Wallis test with Dunn’s multiple comparisons). Right graph shows median, interquartile range and minimum to maximum values of at least three independent experiments. ns, non-significant (one-way Anova with Dunnett’s multiple comparisons). In both graphs each data point represents one cell. (**F**) Quantification of proliferating T cells after Centrinone treatment according to CFSE labeling. DCs were treated with the indicated concentrations of Centrinone and loaded with or w/o antigen. Graph displays mean values ± s.d. Each data point represents one independent experiment with at least *N* = 10.000 cells analyzed per condition. Cells were derived from three different mice. *, *P* < 0.0332; **, *P* < 0.0021 (two-way Anova with Dunnett’s multiple comparisons).

To specifically deplete centrosomes in DCs, we treated BMDCs with the polo-like kinase 4 (PLK4) inhibitor Centrinone during differentiation and maturation to inhibit the formation of new daughter centrioles (Wong *et al*, 2015). Centrinone treatment did not interfere with terminal differentiation of DCs as determined by upregulation of DC-specific cell surface markers such as CD11c and MHCII (**Fig. EV1E**) but efficiently depleted CETN2-GFP^+^ foci in more than 60% of all mature cells (**Fig. 1B**). Note that Centrinone-treatment solely inhibits the generation of new procentrioles but does not deplete existing centrioles leading to a mixed population of cells that contain either no or one centriole, two, three or four, or more than four centrioles (**Fig. 1B**, sketch). When probing cells with antibodies against γ-tubulin to monitor PCM proteins surrounding the centrioles, we found residual staining present in Centrinone-treated cells, which appeared less intense than in untreated controls (**Fig. 1C**). This indicates some degree of residual PCM organization after Centrinone treatment. To confirm lack of centrioles at higher resolution, we carried out expansion microscopy of Centrinone-treated and control cells. Mature DCs were immobilized on poly-L-lysin coated cover slips and stained against acetylated (ac)-tubulin to identify centrioles, γ-tubulin to monitor PCM proteins and N-Hydroxysuccinimide (NHS) ester to unspecifically label free amino groups of proteins. In control samples, we routinely found cells with two, four and more than four barrel-shaped centrioles identified by ac-tubulin and NHS ester staining, that were surrounded by PCM proteins (**Fig. 1D**, left). In the presence of PLK4-inhibition, ac-tubulin staining was not present, but cells had pronounced PCM structures shown by the presence of γ-tubulin (**Fig. 1D**, right). Similarly, NHS ester staining did not yield prominent barrel-shaped structures that co-localized with ac-tubulin staining in control cells. These results indicate that pharmacological inhibition of PLK4 leads to depletion of centrioles but maintains part of the centrosomal PCM in DCs. In line with these findings, residual PCM organization has recently been described in cytotoxic T lymphocytes after genetic depletion of centrioles (Tamzalit *et al*, 2020).

The presence of γ-tubulin in the absence of centrioles prompted us to further examine whether MT architecture was affected by lack of centrioles. Immunostaining of Centrinone-treated and control cells against α-tubulin revealed that acentriolar cells exhibited the same number of MTs and MTOCs than control cells demonstrating that MT integrity was unperturbed in cells lacking centrioles (**Fig. 1E**). When co-culturing antigen-loaded Centrinone-treated cells with OT-II specific T cells, we found that cells lacking centrioles exhibited a significantly diminished capacity for CD4^+^ T cell priming compared to control cells (**Fig. 1F**). As MT organization was not affected by loss of centrioles, these results reveal that centrioles and PCM integrity in APCs are essential for efficient T cell activation independent of centrosomal MTOC function.

To further analyze the role of centrosomal MT nucleation during T cell activation, we next perturbed MT growth in DCs by pre-treating cells with the MT destabilizing agent pretubulysin (Ullrich *et al*, 2009; Braig *et al*, 2014) (**Fig. 2A**). Pretubulysin treatment efficiently depolymerized MT filaments in DCs (**Fig. 2B,C**). Importantly, wash-out of pretubulysin did not result in full recovery of MTs even after 24h of wash-out thus allowing to study the role of MTs specifically in DCs, while leaving the T cell’s MT cytoskeleton unaffected (**Fig. 2D,E**). To this end, we first loaded mature DCs with OVAp and subsequently treated cells for 1h with pretubulysin to disassemble MT filaments after antigen loading. Pre-treated and control DCs were either washed two times (wash-out) or directly (w/o wash-out) co-cultured with OT-II-specific T cells. We found that T cell activation was markedly reduced in the presence of pretubulysin (**Fig. 2F**). Notably, also after drug wash-out, DCs displayed a substantially reduced T cell activation capacity that decreased from 62% activated T cells in control samples to 27% after pretubulysin treatment and wash-out in the presence of 0.1 *μ*g/ml OVAp (**Fig. 2F, right**). These results highlight that centrosomal MT growth in DCs is an immediate prerequisite for efficient T cell activation. In summary, our results emphasize the importance of an intact centrosome and centrosomal MT array in APCs for eliciting CD4^+^ T cell responses.

**Figure 2.**
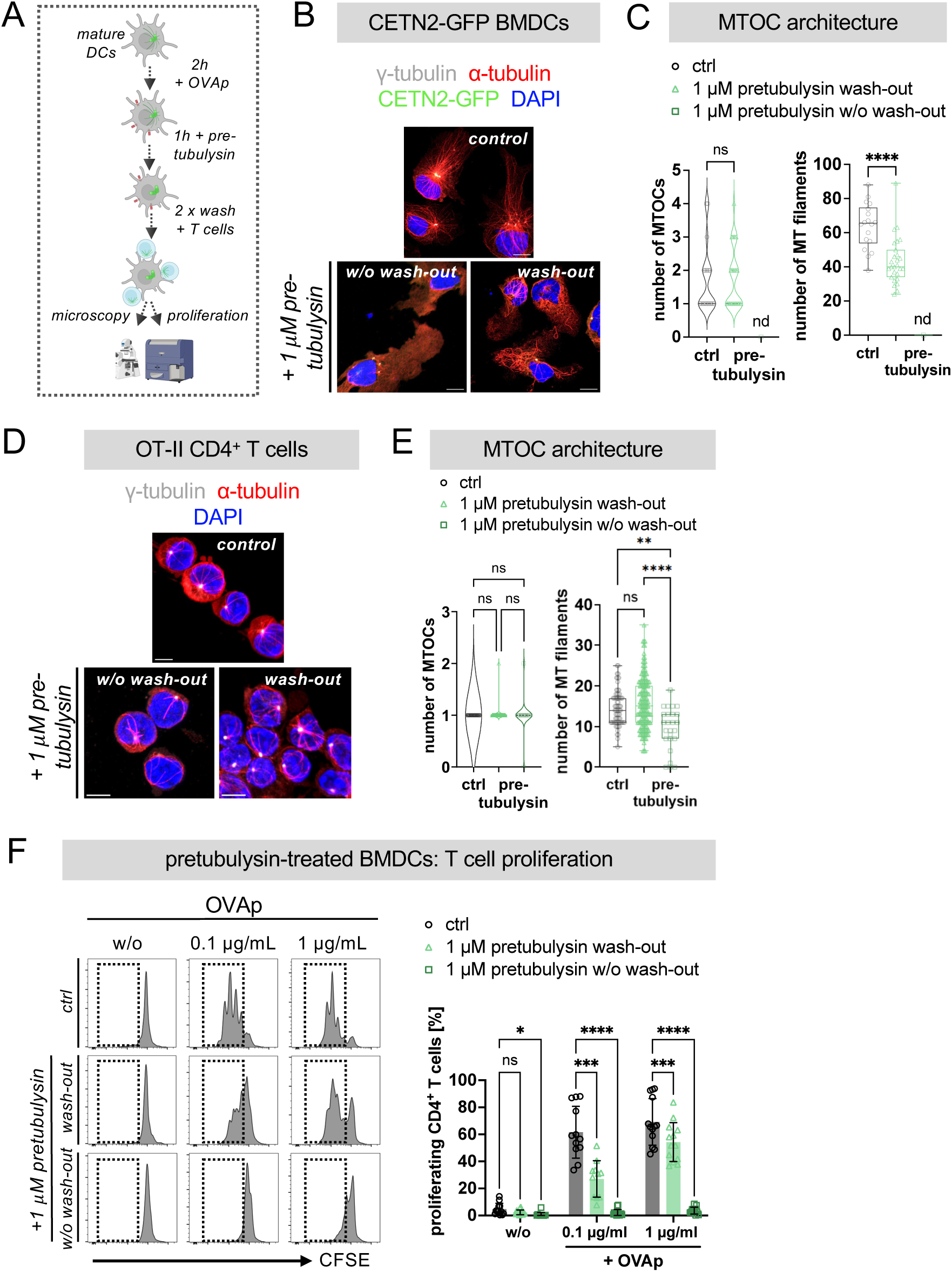
An intact MT cytoskeleton in APCs is required for efficient T cell activation. (**A**) Schematic representation of experimental workflow: CETN2-GFP expressing BMDCs were loaded with OVAp for 2h and subsequently treated with 1 µM pretubulysin. After drug wash-out or without wash-out, T cells were added and cell-cell conjugates analysed via confocal microscopy and flow cytometry. Picture created with BioRender. (**B**) Immunostaining of CETN2-GFP expressing BMDCs after pretubulysin treatment and confinement under agarose. Merged channels of CETN2-GFP (green), α-tubulin (red), γ-tubulin (white) and DAPI (blue) are shown. Scale bars, 10 *μ*m. (**C**) Quantification of MTOCs (left) and MT filaments (right) in BMDCs treated with pretubulysin. Left graph shows median and distribution of data points of two independent experiments (Mann-Whitney test). Right graph shows median, interquartile range and minimum to maximum values of two independent experiments. Each data point represents one cell. ****, P < 0.0001 (Mann-Whitney test). nd, not determined. (**D**) Immunostaining of T cells co-cultured for 24 h with BMDCs previously treated with pretubulysin according to (A). Merged channels of α-tubulin (red), γ-tubulin (white) and DAPI (blue) are shown. Scale bars, 10 µm. (**E**) Quantification of MTOCs and MT filaments in T cells from (D). Left graph shows median and distribution of data points of two independent experiments. ns, non-significant. (Kruskal-Wallis test with Dunn’s multiple comparisons). Right graph shows median, interquartile range and minimum to maximum values of two independent experiments. Each data point represents one cell. **, *P* < 0.021; ****, *P* < 0.0002 (one-way Anova with Dunnett’s multiple comparisons). (**F**) Quantification of T cell proliferation after co-culture with pretubulysin-treated BMDCs. T cell proliferation was assessed in the absence (wash-out) or presence (no wash-out) of pretubulysin. Graph displays mean values ± s.d. Each data point represents one independent experiment with at least *N* = 10.000 cells analyzed per condition. Cells were derived from three different mice. *, *P* < 0.0332; ***, *P* < 0.0002; ****, *P* < 0.0001 (two-way Anova with Dunnett’s multiple comparison). ns, non-significant.

### Enhanced MTOC activity in DCs with amplified centrosomes during IS formation

Our data highlight a crucial role of centrioles and centrosomal MT filaments for efficient T cell activation. While centrosome numbers are generally limited to one in G1 phase and two prior to mitosis, DCs amplify centrioles upon antigen encounter (Weier *et al*, 2022). To address whether extra centrioles in DCs impact T cell activation, we visualized centrosomes and MT arrays during DC-T cell interaction. As extra centrioles in DCs arise due to mitotic defects, which are accompanied by tetraploidization, we focused on diploid cells and first examined the functionality of extra centrioles in APCs during DC-T cell interaction in fixed samples. 82% of mature BMDCs showed a 2N DNA profile of which 25%±8% of cells contained ≥3 centrioles (**Figs. 1A** and **3A**). We then mixed OVAp loaded CETN2-GFP expressing DCs with OT-II T cells and fixed cell-cell conjugates after 2h of conjunction. Cells were immuno-stained against *γ*-tubulin to monitor PCM proteins surrounding the centrioles and *α*-tubulin to visualize MT filaments (**Fig. 3B**). We next determined the levels of PCM proteins in cells with one and amplified centrioles during DC-T cell contacts. We found that levels of *γ*-tubulin were increased in cells with extra centrioles indicating that amplified centrioles are able to recruit PCM (**Fig. 3C**). We obtained similar results in the absence of T cells suggesting that enhanced PCM recruitment in DCs with amplified centrioles is a cell intrinsic property and independent of DC-T cell contact formation (**Figs. 3D** and **EV2A-C**). High-resolution deconvolution microscopy further allowed us to identify MT filaments growing from individual MTOCs and quantify filament numbers within defined areas around the centrosome. According to the role of *γ*-tubulin in promoting MT nucleation, we found increased numbers of MT filaments in the presence of extra centrioles during antigen-specific DC-T cell interactions (**Fig. 3E**). These results demonstrate that centrosomal MT nucleation capacity in DCs is increased in the presence of additional centrioles leading to over-active MTOCs and a larger number of cytoplasmic MT filaments.

**Figure 3.**
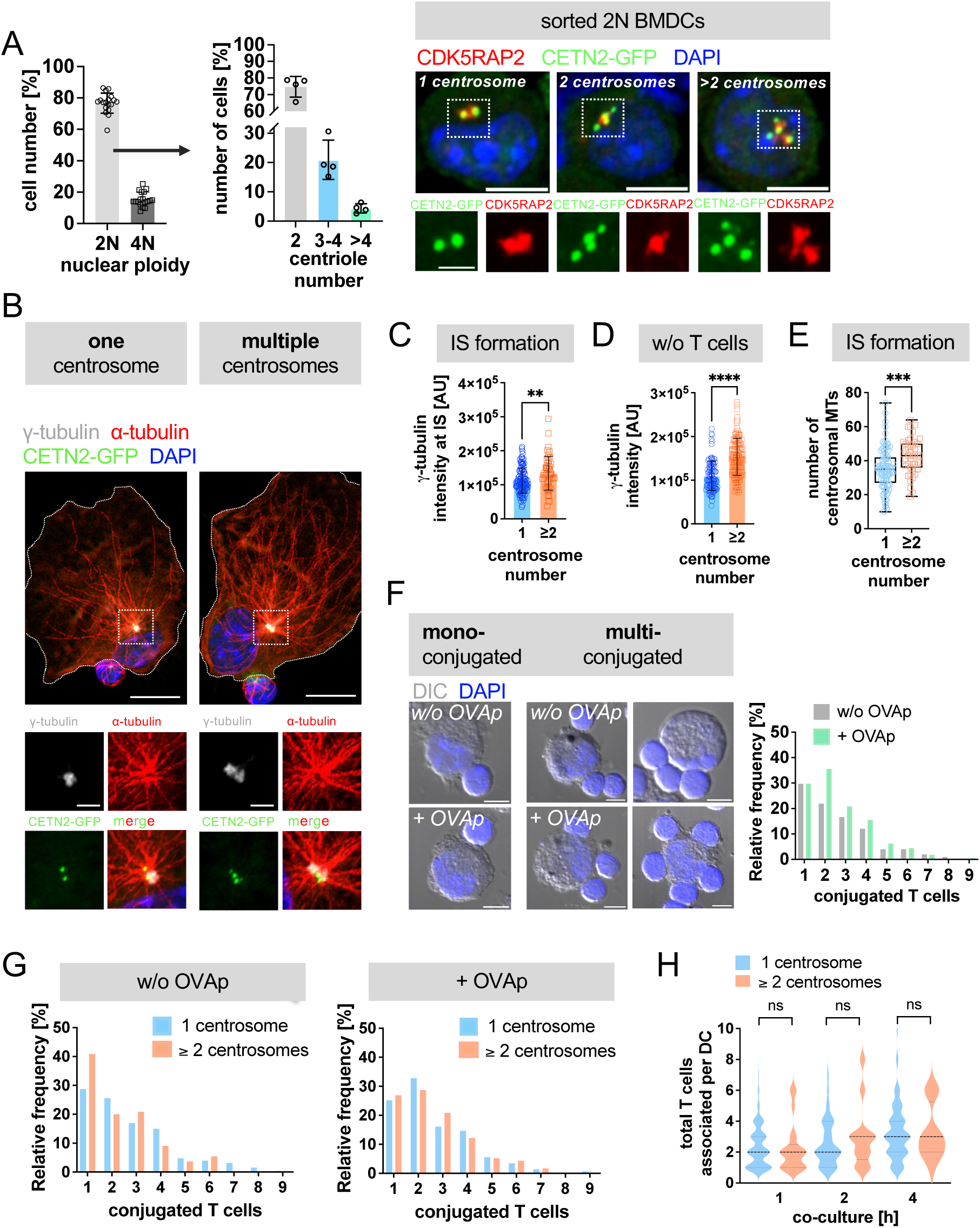
IS formation in the presence of multiple centrosomes. (**A**) Left: quantification of 2N and 4N cells according to DNA content (*see also* Fig. 1A). Middle: quantification of centrosome numbers in sorted mature 2N CETN2-GFP BMDCs according to CETN2-GFP/γ-tubulin^+^ foci. Graph shows mean values ± s.d. of 4 independent experiments with at least *N* = 200 cells analyzed per experiment. Right: immunostaining of centrioles and PCM in sorted 2N mature CETN2-GFP BMDCs. Merged channels of CDK5RAP2 (red), CETN2-GFP (green) and DAPI (blue) are shown. Scale bars, 5 *μ*m. Insets show magnification of indicated regions. Individual channels of CETN2-GFP (green) and CDK5RAP2 (red) are shown. Scale bars, 2 *μ*m. (**B**) Immunostaining of MT filaments in sorted 2N mature CETN2-GFP expressing BMDCs loaded with OVAp and forming conjugates with T cells. White dotted box indicates magnified region below. Merged and individual channels of CETN2-GFP (green), γ-tubulin (white), α-tubulin (red) and DAPI (blue) are shown. Scale bars, 10 *μ*m (upper panels) and 2 *μ*m (insets bottom). (**C** and **D**) Quantification of γ-tubulin signal intensities at the centrosome in DCs forming conjugates with T cells (**C**) and in the absence of T cells **(D**). Graphs show mean values ± s.d. of one out of three independent experiments. Each data point represents one cell. **, *P* < 0.0021; ****, *P* < 0.0001 (Mann-Whitney test). (**E**) Quantification of MT numbers emanating from defined regions around centrosomes in sorted 2N CETN2-GFP BMDCs loaded with OVAp and forming contacts with T cells. Graph displays median, interquartile range and minimum to maximum values. Each data point represents one cell derived from 3 independent experiments. ***, *P* < 0.0002 (two-tailed, unpaired Student’s *t*-test). (**F**) Left: merged channels of differential interference contrast (DIC, gey) and DAPI (blue) of DC-T cell conjugates in the absence (upper panels) and presence of OVAp (lower panels). Scale bars, 5 *μ*m. Right: quantification of frequency distribution of bound T cells per DC. Graphs display normalized values ± s.d. *N* > 200 cells analyzed per condition. (**G**) Histogram of frequency distribution of bound T cells to a single DC after 2h of conjugate formation in cells with one (blue) and ≥2 (red) centrosomes. *N* > 100 cells analyzed per condition. (**H**) Quantification of bound T cells per DC after 1, 2 and 4h of conjugate formation. Graph shows median and distribution of values of 5 independent experiments (Kruskal-Wallis test with Dunn’s multiple comparisons). *N* = 132/59/97 cells analyzed (1h/2h/4h). ns, non-significant.

### DCs form multiple T cell contacts independently of centrosome numbers

To address whether enhanced centrosomal MT nucleation in DCs with amplified centrosomes correlates with a higher capacity to form multiple T cell contacts, we determined the frequency distribution of bound T cells in relation to DC centrosome numbers. In the presence of excess T cells (1:5 DC/T cell ratio), DCs engaged with either one T cell (mono-conjugated) or several T cells simultaneously (multi-conjugated) (**Fig. 3F**). Cell-cell contacts were also formed in the absence of OVAp as described before (Benvenuti *et al*, 2004; Mittelbrunn *et al*, 2009). As CD4^+^ T helper synapses comprise at least four distinct stages that proceed over several hours and are associated with morphological shape changes as well as reorganization of cytoskeletal components in both cell types (Ueda *et al*, 2011), we analysed conjugate formation after 1, 2 and 4h after conjunction. Cells carrying a single centrosome bound on average two to three T cells simultaneously in the presence of excess T cells (**Fig. 3G**). We did not detect significant differences in the number of interacting T cells between DCs with one or multiple centrosomes at distinct time points of conjugation (**Fig. 3G,H**). Thus, we concluded that the capacity to form cell-cell contacts between DCs and T cells occurs independently of centrosome numbers and the levels of centrosomal MT nucleation in DCs.

### Enhanced T cell activation in the presence of amplified centrosomes in DCs

To further address whether the presence of additional centrosomes impacts T cell activation, we sought to simultaneously visualize T cell activation and centrosome numbers. To this end, we made use of Nur77^GFP^ transgenic mice, which express GFP under the control of the Nur77 promotor (Liu *et al*, 1994). Nur77 gene expression is up-regulated by antigen-dependent TCR stimulation but not by other homeostatic or inflammatory signals (Moran *et al*, 2011). Accordingly, in this mouse model, the levels of GFP expression directly reflect TCR signal strength and reach a maximum expression after 12-24h of TCR stimulation (Liu *et al*, 1994). To estimate the timing of T cell activation after DC encounter in the context of MHCII antigen presentation, we crossed Nur77^GFP^ mice with OT-II transgenic mice. Flow cytometric analysis revealed that Nur77-dependent GFP expression in T cells was generally low in the absence of antigen and after 1h of co-culture with OVAp-loaded CETN2-GFP expressing DCs, while 2h after mixing a clear GFP signal appeared, which strongly increased after 4h, 6h and 20h of incubation (**Fig. 4A,B**). At 20h of co-culture in the presence of antigen, 67% of all T cells showed a prominent GFP-signal indicating efficient activation. By contrast, Nur77^GFP^ levels remained low in the absence of antigen at all time points analyzed demonstrating antigen-dependent T cell activation (**Fig. 4B**).

**Figure 4.**
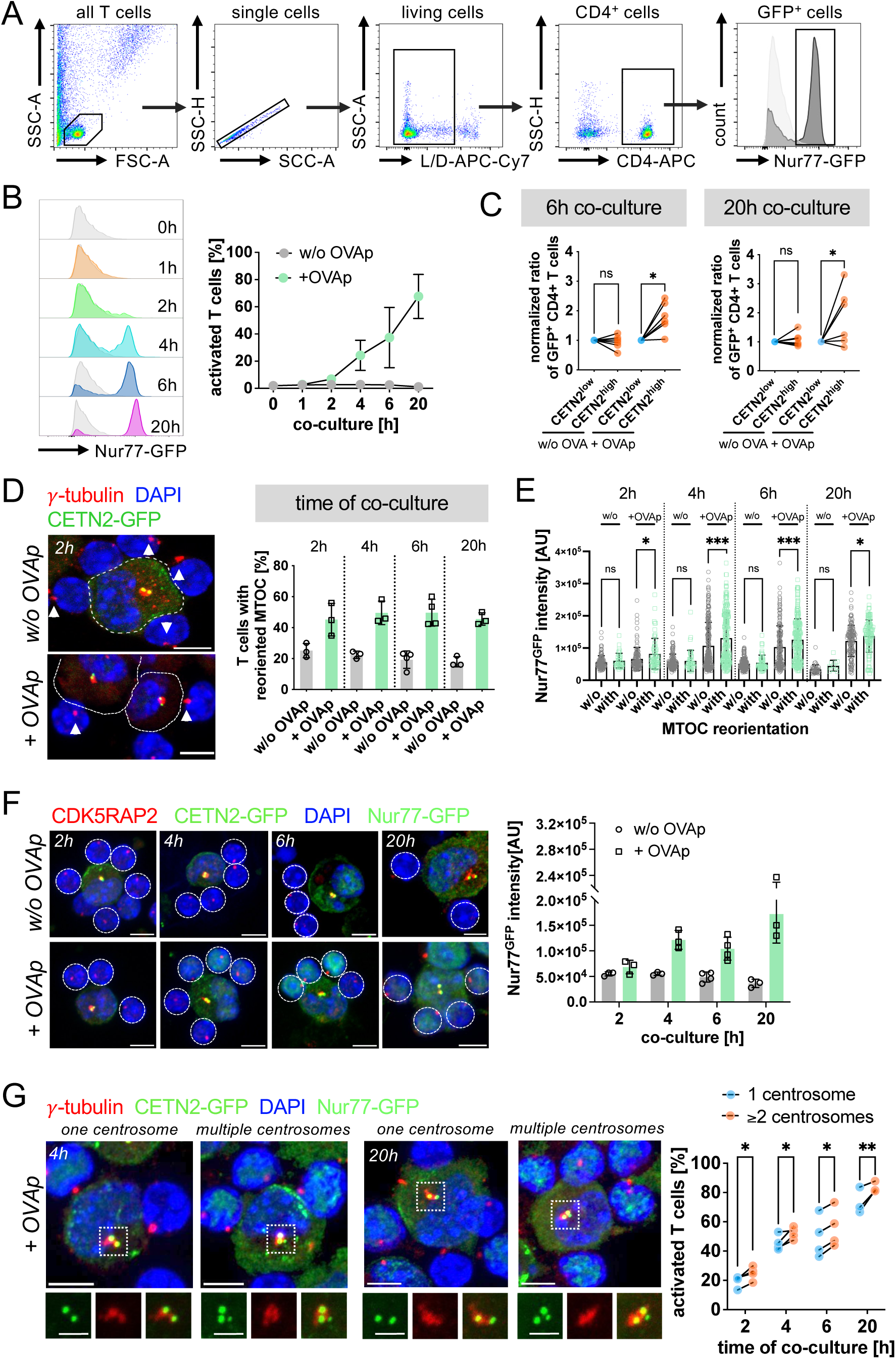
Enhanced T cell activation in the presence of multiple centrosomes. (**A**) Gating strategy for analyzing Nur77^GFP^ expression in CD4^+^ T cells. One representative experiment out of 4 is shown. T cells without DC co-culture served as control and were included as light grey filled line. (**B**) Histogram (left) and quantification of Nur77^GFP^ expression (right) after different time points of DC-T cell co-culture in the presence (+ OVAp) or absence (w/o OVAp) of OVAp. Graph represents mean values ± s.d. of 4 independent experiments. *N* = 10.000 cells analyzed per condition. (**C**) Normalized ratio of GFP^+^/CD4^+^ T cells after 6 and 20h of co-culture with OVAp-pulsed 2N BMDC subpopulations. Each data point represents one independent experiment with pairing between sorted 2N CETN2-GFP^low^ and CETN2-GFP^high^ expressing cells. CETN2-GFP^low^ values were normalized to 1. *, *P* < 0.0332 (two-tailed, paired Student’s *t*-test). (**D**) Left: Immunostaining of γ-tubulin (red) in DC-T cell conjugates. Merged channels of CETN2-GFP (green), γ-tubulin (red) and DAPI (blue) are shown. DC outline is indicated with dotted line. White arrowheads point to the T cell’s centrosome. Scale bars, 5*μ*m. Right: quantification of MTOC polarization towards the IS in T cells after different timepoints of co-culture. Graph displays mean values ± s.d. Each data point represents one independent experiment with *N* > 100 cells analyzed per condition. (**E**) Quantification of GFP signal intensity in T cells in the presence and absence of antigen and in dependence of MTOC reorientation in the T cell towards the IS. Graph shows one representative experiment out of at least three independent experiments. *, *P* < 0.0332; ***, *P* < 0.0021 (one-way Anova with Kruskal-Wallis multiple comparisons). (**F**) Visualization and quantification of Nur77^GFP^ expression in DC-T cell conjugates after different timepoints of co-culture. Left: immunostaining of CDK5RAP2 (red) in CETN2-GFP (green) BMDCs after co-culture with Nur77^GFP^/OT-II CD4^+^ T cells. Nuclei were counterstained with DAPI (blue). Dotted circles indicate areas of GFP measurements. Scale bars, 5 *μ*m. Right: quantification of GFP signal intensity in T cells in the presence and absence of antigen. Graph displays mean values ± s.d. Each data point represents one independent experiment with N >100 cells analyzed per experiment. (**G**) Analysis of T cell activation in dependence of centrosome numbers. Left: immunostaining of γ-tubulin in DC-T cell conjugates. Merged and individual channels of CETN2-GFP (green), Nur77^GFP^ (green), γ-tubulin (red) and DAPI (blue) are shown. Indicated regions are magnified and shown below. Scale bars, 5 *μ*m (top). Scale bars, 2 µm (magnified region). Right: quantification of T cell activation according to Nur77^GFP^ signal intensities in dependence of DC centrosome numbers for different time points of DC-T cell co-culture. Each data point represents one independent experiment with pairing between cells with one (blue) and multiple (red) centrosomes from one experiment. N > 100 cells analyzed per condition. *, *P* < 0.0332; **, *P* < 0.0021 (two-tailed, paired Student’s *t*-test). ns, non-significant.

To elucidate whether T cell activation changes in the presence of different centrosome numbers, we first sorted BMDCs according to CETN2-GFP signal intensities, which leads to two DC subpopulations enriched for either one centrosome (CETN2-GFP low) and cells with two and more centrosomes (CETN2-GFP high) ((Weier *et al*, 2022), and **Fig. EV3A,B**). The percentage of cells carrying multiple centrosomes ranged from 13–26% in the CETN-GFP low population and 30–49% within the CETN2-GFP high population, leading to an average enrichment of cells with multiple centrosomes by a factor of 2.0 (**Fig. EV3C**). Sorted cells were further loaded with OVAp and incubated for either 6h or 20h with Nur77^GFP^/OT-II transgenic T cells. After both time points of co-culture, the number of Nur77^GFP^-positive cells was significantly higher when T cell priming was accomplished with DCs enriched for multiple centrosomes indicating that these cells activate a larger number of T cells within the same time period compared to cells with only a single centrosome (**Fig. 4C**).

To directly observe T cell activation on a single cell level and in relation to centrosome numbers, we immobilized DC-T cell conjugates at distinct time points of co-culture and determined integrated fluorescence intensities of GFP levels in T cells. To distinguish between distinct numbers of centrosomes, we used CETN2-GFP expressing DCs and counterstained centrosomes with antibodies against γ-tubulin or CDK5RAP2 to simultaneously allow visualization of the T cell MTOC. We observed polarization of the T cell’s centrosome to the nascent IS at all time points analyzed, indicating efficient IS formation (**Fig. 4D**). Similar to our flow cytometric analysis, GFP levels were significantly higher in the presence of antigen and steadily increased from 2h to 6h of co-culture demonstrating TCR activation and antigen-specific T cell priming (**Fig. 4F**). Nur77-dependent GFP expression was elevated in T cells that had already reoriented its centrosome compared to cells with a centrally localized MTOC further demonstrating that centrosome reorientation in T cells correlates with T cell activation (**Fig. 4E**). We next determined T cell activation in relation to the number of centrosomes present in the conjugated DC. To this end, we defined a threshold for T cell activation based on the frequency distribution of GFP signal intensities in T cells co-cultured in the absence of antigen (**Fig. EV3D,E**). We found that as early as 2h of co-culture, the number of GFP^+^ T cells was elevated after priming with DCs that contain amplified centrosomes and further increased after 4, 6 and 20h of mixing (**Fig. 4G**). In accordance to our flow cytometric analysis, these results demonstrate that the total number of activated T cells is higher when the APC contains amplified centrosomes. Altogether, our findings provide evidence that cells with extra centrosomes activate T cells faster compared to cells with only a single centrosome confirming optimized T cell responses in the presence of amplified centrosomes in APCs.

### Multiple centrosomes cluster and localize close to the cell center in DCs during IS formation

Centrosome polarization has been reported in DCs during the interaction with naïve CD8^+^ T cells. MTOC reorientation in DCs depends on the Rho GTPase Cdc42 and enables targeted delivery of vesicles containing T cell stimulatory molecules (Pulecio *et al*, 2010). Based on these observations, we reasoned that one possible explanation for the observed accelerated T cell priming capacity in the presence of multi-numerous centrosomes could be de-clustering of extra centrosomes, which subsequently move and reorient to distinct contact sites in order to efficiently deliver stimulatory molecules to the respective IS (**Fig. 5A**). To test this hypothesis, we measured intracentrosomal distances (between pairs of centrioles) and average distances between all centrioles in DCs with one and multiple centrosomes after conjugate formation in the presence or absence of OVAp. We further distinguished whether cells form mono- or multi-conjugated synapses (**Fig. 5B,C**). Analysis of intracentrosomal distances in cells with a single centrosome treated with or without OVAp for distinct time points of co-culture (1, 2, 4h) revealed distances of 0.8-1.1 *μ*m between individual centrioles independently of the number of T cells attached (**Fig. 5B**). Similarly, average distances in cells with multiple centrosomes ranged between 0.9-1.3 *μ*m and did not show prominent differences between OVAp loaded and unloaded cells suggesting that multiple centrosomes congregate together and cluster during antigen-specific DC-T cell contacts. Average distances were indistinguishable not only when cells formed single contacts but also in multi-conjugated cells (**Fig. 5C**).

**Figure 5.**
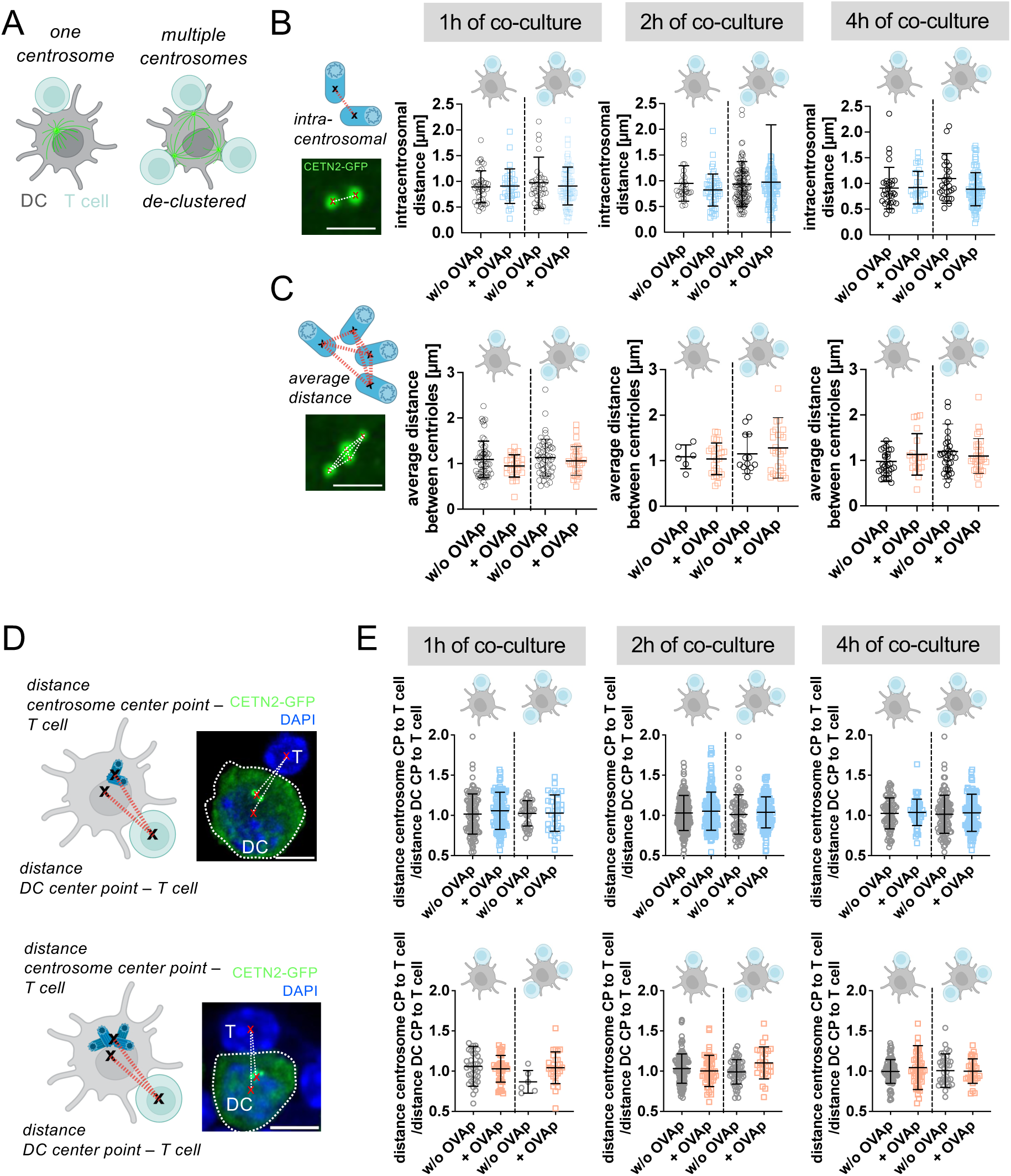
Centrosome configuration during antigen-specific DC-T cell contacts. (**A**) Sketch illustrating centrosome de-clustering and centrosome polarization in DCs toward the IS in the presence of one (left) and multiple centrosomes (right). Picture in B and C created with BioRender. Quantification of intracentrosomal distances (**B**) and average distances (**C**) in DCs with one and multiple centrosome(s) at different time points of co-culture and bound to one or several T cells (separated by dashed line and indicated on top). Graphs display mean values ± s.d. Each data point represents one cell derived from 5 independent experiments. Sketches and pictures indicate centrosome configuration (CETN2-GFP) in DC-T cell contacts. Scale bars, 2 *μ*m. (**D**) Sketches and confocal CETN2-GFP images indicating distances between centrosome center point (CP) and T cell CP, and DC CP and T cell CP in cells with one (upper) and multiple (lower) centrosomes. Nuclei were counterstained with DAPI (blue). Scale bars, 5 *μ*m. (**E**) Quantification of ratio between distance from centrosome CP to T cell CP and distance DC CP to T cell in cells with one (upper row; blue) and multiple (lower row, red) centrosomes. Graphs show mean values ± s.d. for DCs attached to one T cell (left) and multiple T cells (right). Each data point represents on cell derived from 5 independent experiments.

To further analyze centrosome positioning in DCs, we quantified the ratio of distances between the DC centrosome and the T cell and between the DC center point and T cells in the presence or absence of antigen (**Fig. 5D**). Our analysis revealed that 4h after co-culture, the centrosome in DCs was still located in close proximity to the cell center and the nucleus (**Figs. 5E** and **EV4A**). This behavior was not only observed in multi-conjugated cells, but also when single synapses were formed. These results suggest that MTOC positioning in DCs was independent of the number of centrosomes and T cells conjugated. Moreover, our findings demonstrate that MTOC polarization toward the IS in DCs is dispensable for T cell activation.

To further study DC centrosome dynamics with high spatio-temporal resolution in particular during early DC-T cell encounters, we recorded time-lapse images of DC-T cell conjugates. To this aim, CETN2-GFP expressing DCs were loaded with OVAp, mixed with OVA-specific OT-II T cells and imaged directly after mixing. To monitor efficient DC-T cell interaction, we visualized intracellular calcium (Ca^2+^) influx in T cells and considered only interactions with long-lasting Ca^2+^ responses. In cells containing a single centrosome, intracentrosomal distances did not increase upon contacts with one or multiple T cells (**Fig. 6A** and **movies EV1** and **EV2**). Similarly, average distances in cells with amplified centrosomes remained unaltered in conjunction with one or multiple T cells (**Fig. 6B** and **movies EV3** and **EV4**). Moreover, no net movement of single or multiple centrosome(s) in DCs could be observed within the imaging period (0-1h after conjugation) in the presence or absence of antigen (**Fig. 6C,D** and **movies EV1-4**). Instead, centrosomes were found in close proximity to the nucleus independently of the number of T cells bound, similar to our fixed samples (**Fig. EV4A,B**). Overall, our results demonstrate that amplified centrosomes cluster during IS formation and collectively stay close to the cell center when mono- or multi-conjugated contacts are formed.

**Figure 6.**
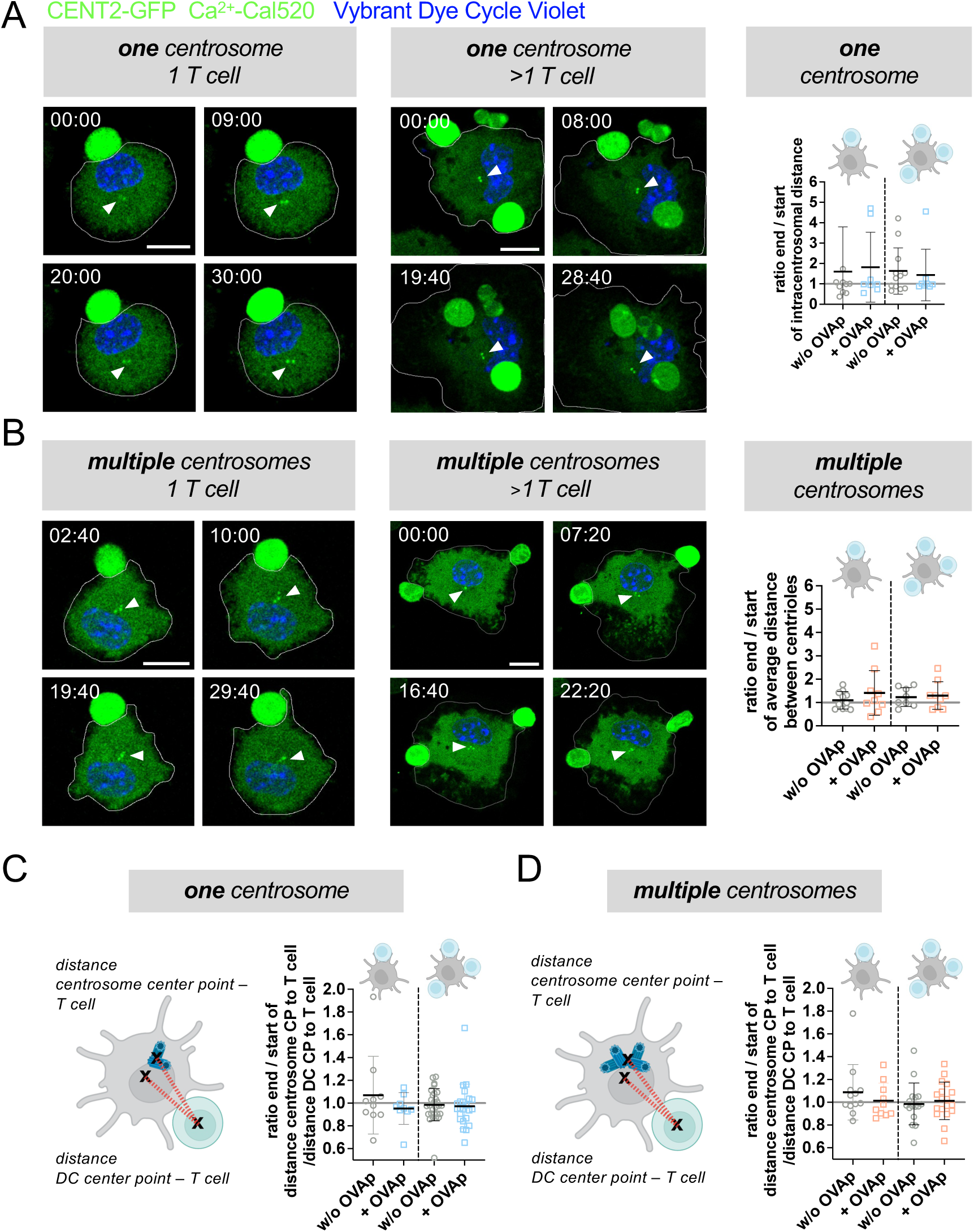
Live cell imaging of centrosome dynamics at the IS. (**A+B**) Time-lapse live-cell confocal microscopy of antigen-specific DC-T cell contacts. Left: merged images of CETN2-GFP (green), Ca^2+^-Cal520 (green) and DNA stain (Vybrant Dye Cycle Violet, blue) are shown. Frames were collected every 20 s. Only selected frames are shown in montage with precise time points indicated in min:sec. White arrow heads point to centrosome(s) position. Scale bars, 10 *μ*m. (**A**) Right: quantification of ratio of end vs. start intracentrosomal distance in cells with only a single centrosome (left, blue) and dividing mono-conjugated (left half) and multi-conjugated synapses (right half). Graphs show mean values ± s.d. Each data point represents one cell recorded for at least 30-60 minutes from at least 5 independent experiments. (**B**) Right: quantification of ratio of end vs. start average distance between centrioles in cells with multiple centrosomes and dividing mono-conjugated (left half) and multi-conjugated synapses (right half). Graph shows mean values ± s.d.. Each data point represents one cell recorded for 30-60 minutes from minimum 5 independent experiments. (**C**) Left: sketch indicating distances between centrosome CP to T cell CP, and DC CP to T cell CP in cells with one centrosomes. Right: quantification of ratio of end vs. start of distance centrosome CP to T cell CP normalized to the movement of the DC. Graph shows mean values ± s.d. Each data point represents one cell recorded for 30-60 minutes from at least 5 independent experiments. (**D**) Left: sketches indicating distances between CP centrosome CP to T cell CP, and DC CP to T cell CP in cells with multiple centrosomes. Right: quantification of ratio of distances between the centrosome CP to T cell at the end vs. beginning of imaging (normalized to the movement of the DC). Graph represents mean values ± s.d. Each data point represents one cell derived from 5 independent experiments.

### De-clustering agents impair T cell activation

As we observed centrosome clustering during DC-T cell contacts, we next addressed the importance of this phenomenon for the induction of T cell activation. Therefore, DCs were treated with the de-clustering agent PJ-34, a poly-ADP-ribose polymerase inhibitor with no MT binding activity in proliferating cells (Chiarugi *et al*, 2003; Castiel *et al*, 2011). PJ-34 was administered for 3h on mature DCs to test the effect on centrosome organization. Under control conditions (H_2_O), cells containing a single centrosome display mean intracentrosomal distances of 0.7±0.3 *μ*m, while intercentrosomal distances in cells with multiple centrosomes were slightly larger with average distances of 1.3±0.7 *μ*m (**Fig. 7A,B**). PJ-34 treatment lead to a significant shift to larger distances in particular of intercentrosomal distances in cells carrying multiple centrosomes (2.2±1.6 *μ*m) indicating that pairs of centrioles get dispersed (**Figs. 7B** and **EV4C**). Immunostaining against *α*-tubulin and analysis of MT filaments emanating from either clustered or dispersed centrosomes revealed that PJ-34-treated cells nucleate slightly more MT filaments compared to control cells (**Fig. 7C**). Intriguingly, when co-cultering either PJ-34-treated or control OVAp-loaded DCs with OT-II T cells, we found that in the presence of de-clustered centrosomes, T cell activation was significantly diminished compared to control cells (**Fig. 7D,E**) demonstrating that de-clustering of extra centrosomes impairs T cell activation. In summary, our findings highlight a crucial role for proper centrosome arrangement in APCs to efficiently activate T cells.

**Figure 7.**
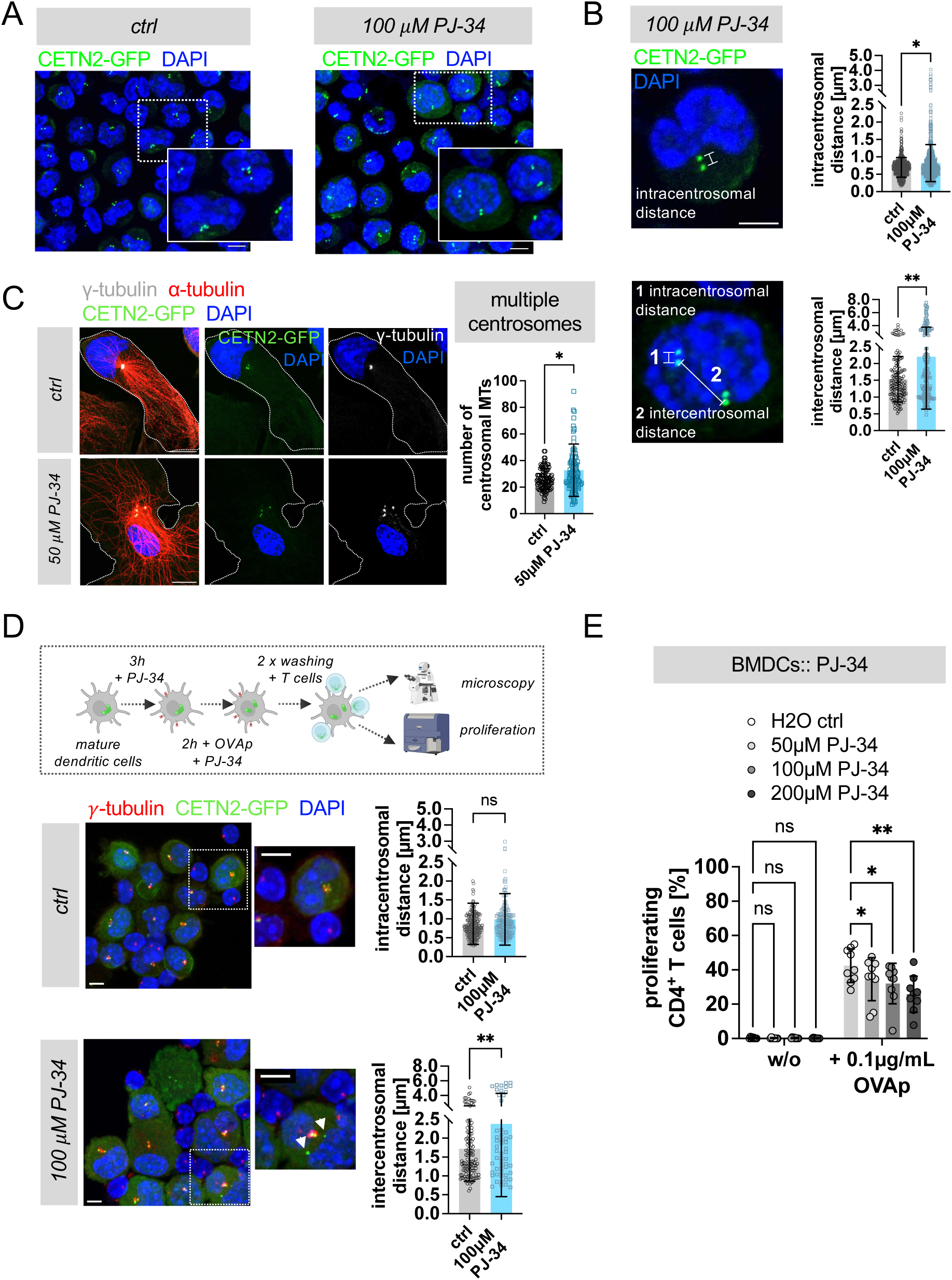
Centrosome de-clustering impairs T cell activation. (**A**) Visualization of centrioles in CETN2-GFP expressing BMDCs after PJ-34 treatment. Insets show magnification of indicated region. Nuclei were counterstained with DAPI (blue). Scale bars, 5 *μ*m. (**B**) Visualization and quantification of intracentrosomal (upper panel) and intercentrosomal (lower panel) distances in cells treated with PJ-34. Merged images of CETN2-GFP (green) and DAPI (blue). Scale bars, 2 *μ*m. Graphs display mean values ± s.d. Scale bars, 2 *μ*m. Each data point represents one cell derived from three independent experiments (two-tailed, unpaired Student’s *t*-test with Welch’s correction). (**C**) Left: immunostaining of PJ-34-treated and control cells. Merged and individual channels of CETN2-GFP (green), α-tubulin (red), γ-tubulin (white) and DAPI (blue) are shown. Scale bars, 10 *μ*m. Right: quantification of MT numbers emanating from defined regions around centrosomes in 2N PJ-34 treated and control cells. Graph displays mean values ± s.d. Each data point represents one cell derived from two independent experiments. *, *P* < 0.0332 (two-tailed, unpaired Student’s *t*-test). (**D**) Upper panel: schematic experimental layout. Below: Immunostaining of CETN2-GFP (green) expressing DC-T cell conjugates against γ-tubulin (red) after 2h of mixing in the absence or presence of the de-clustering agent PJ-34. Nuclei were counterstained with DAPI (blue). Scale bars, 5 *μ*m. Insets show magnification of indicated regions. Scale bars, 2 µm. White arrowheads point to de-clustered centrioles. Right: Quantification of intracentrosomal and intercentrosomal distances in cells treated with PJ-34. Graphs display mean values ± s.d.. Each data point represents one cell derived from three independent experiments (two-tailed, unpaired Student’s *t*-test). (**E**) T cell proliferation with or without PJ-34 treatment of DCs was quantified by CFSE labelling. Graph displays mean values ± s.d.. Each data point represents one independent experiment with at least N = 10.000 cells analysed per condition and cells derived from three different mice. *, P < 0,0332; **, P < 0.0021; ***, P < 0.0002 (two-way Anova with Dunnett’s multiple comparison). ns, non-significant.

### Modeling T cell priming with geometrically optimal centrosome positioning in DCs

Our experimental data emphasize a crucial role of centrosome positioning close to the cell center in DCs and provide evidence that T cell activation is increased in the presence of amplified clustered centrosomes compared to DCs containing a single centrosome. These observations prompt queries into whether the centrosome’s proximity to the cell center is advantageous for DCs. For instance, a centrosomal position geometrically optimal to the entire cell surface may expedite the delivery of immune signals to the interacting T cells. The optimal position will also provide dynamic MTs faster access to the IS. Therefore, a key task is to examine how MTs achieve optimal centrosome position and the physical consequences of centrosome clustering. In other words, does clustering of centrosomes facilitate efficient MT search for the IS over dispersed centrosomes? And whether the optimal position also corresponds to the mechanically stable position of the centrosomal cluster. To further elaborate on this, we resort to mathematical and computational models to systematically approach these questions. First, we rationalized with simple geometric considerations the physiological consequences of centralized clustered centrosomes during T cell priming. We first seek to explore the geometrically optimal position of the centrosome in DCs relative to the entire cell surface, without explicitly delving into the complexities of MT dynamics. Since the position of the target - the center of the IS on the cell surface - is not predetermined, a centrosomal position with minimal average distance to all possible target points on the cell surface may facilitate efficient delivery of stimulatory molecules to the IS. For cell surface regions directly accessible to the centrosome, the shortest distance is a straight line from the centrosome to the target point on the cell surface. However, for regions no longer visible due to nuclear hindrance, the minimum geometric distance would follow the scheme presented in **Fig. 8A**. This distance corresponds to the path length of MTs on optimal trajectories between the centrosome and the target point on the cell surface. For a detailed calculation, we refer to the *Materials and Methods* section. In the absence of a nucleus, the optimal centrosome position coincides with the geometric center of the cell, as determined through analytical calculations in both circular and spherical cellular geometries and further validated by computational analysis (see *Appendix;* **Fig. EV5A**). However, the presence of a nucleus covering the central region of the cell prevents the centrosome from locating at the cell center. Using computational analysis, we first explore the optimal centrosome position in the presence of a centrally located nucleus and estimate the average geometric distance, < *d_short_* >, as a function of the centrosome’s distance from the cell center, *h_CS_*, or from the nuclear surface *d_CS−NS_*. Our data indicate an optimal perinuclear positioning of the centrosome, marginally shifted from the nuclear surface by < 1 *μm* (**Fig. 8B**). We further investigate the centrosome’s position for various off-centered positions of the nucleus. Note that, for a centrally located nucleus, spherical symmetry of the cell and the nucleus ensures that estimating < *d_short_* > with random target points on the cell surface is independent of the direction of the centrosome placed away from the nucleus. It solely depends on the centrosome’s distance from the nucleus. However, for an off-centered nucleus, < *d_short_* > depends on the specific three-dimensional positioning of the centrosome relative to the nucleus. We discovered that the global minimum of < *d_short_* > is achieved when the centrosome lies on a fixed axis pointed from the nuclear center toward the cell center (shown as Z-axis in the plots; **Fig. EV5B-E**). Therefore, by placing the centrosome at various locations along that axis, we estimate the optimized values of *h_CS_* and *d_CS−NS_*, corresponding to the minimum of < *d_short_* >, and plotted for three different nuclear positions: *d_offset_* = 0 *μm*, representing a centrally located nucleus; *d_offset_* = 6 *μm*, where the cell center is just outside the nuclear surface and *d_offset_* = 12 *μm*, indicating the maximally off-centered nucleus touching the cell surface (**Fig. 8C**). Our findings suggest that positioning the centrosome close to the nuclear periphery is advantageous when the nucleus is at the cell center. The optimal centrosome position shifts toward the cell center as the nucleus becomes off-centered. Overall, our model provides a rational accounting for the geometric positioning of clustered centrosomes near the cell center leading to efficient priming of T cells on the cell surface.

**Figure 8.**
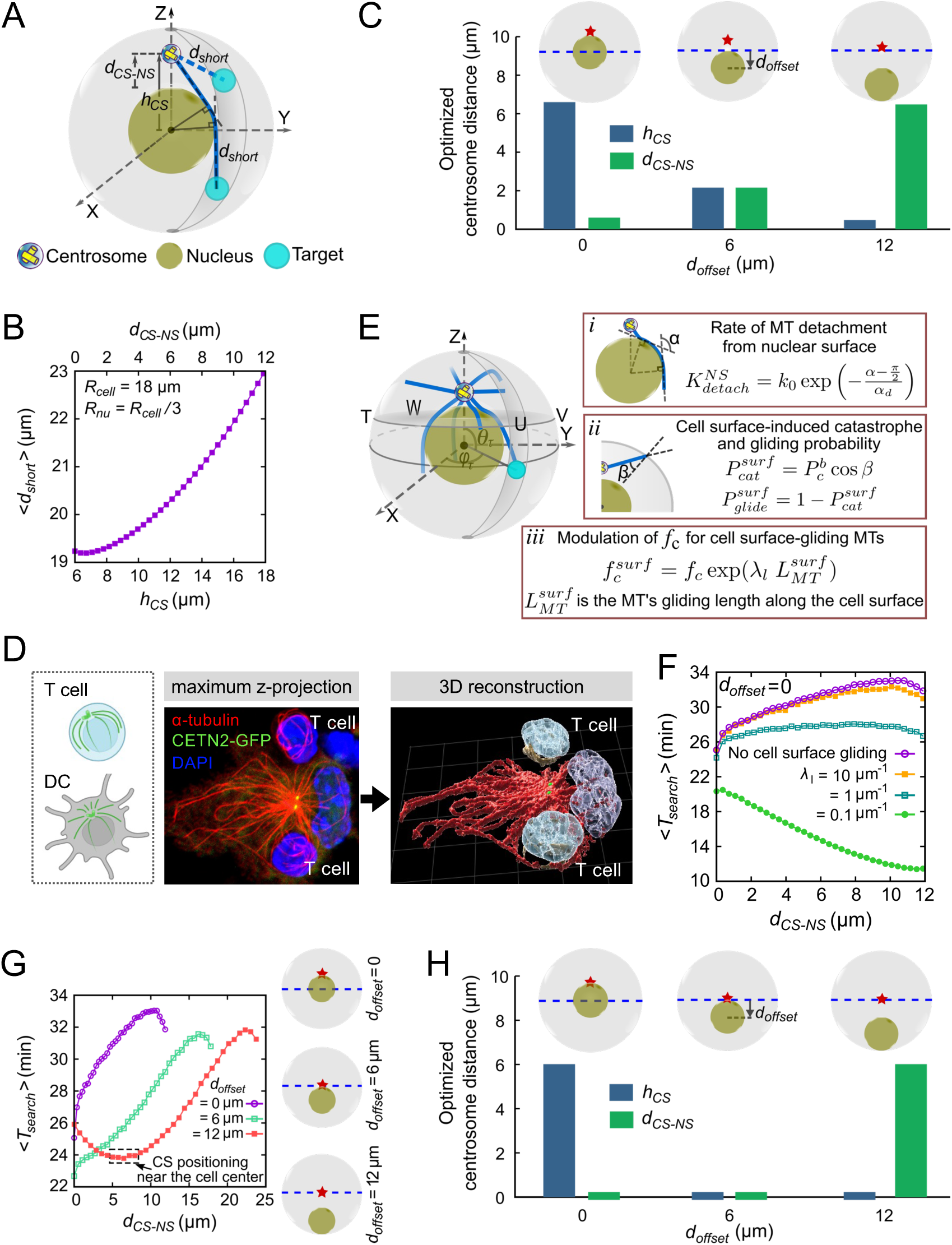
Modeling centrosome positioning in DCs. (**A-C**) Average shortest geometric distance between centrosome and target center on cell surface regulating centrosome position in DCs. (**A**) A schematic representation of the shortest distance, *d_short_*, for the target positions directly or indirectly accessible due to nuclear hindrance. (**B**) Plot of < *d_short_* > vs *h_CS_* in the presence of a nucleus located centrally to the cell. *h_CS_*, represents the centrosome distance from the cell center. The top x-label in (B) represents the centrosome distance away from the nuclear surface, *d_CS−NS_*. (**C**) Optimized values of *h_CS_*and *d_CS−NS_*, corresponding to the minimum of < *d_short_* > for different centrosome positions along the z-axis above the nucleus and for different off-centered positions of the nucleus, described by the values of *d_offset_*. *d_offset_* denotes the distance between the cell center and the nucleus center and is graphically depicted in the inset of (C). (**D**) 3D reconstruction of the MT cytoskeleton in DCs and T cells. Left: sketch illustrating centrosomal MT growth in T cells and DCs. Note that MTs in T cells grow tangentially relative to the cell membrane, while in DCs MTs project astrally from the centrosome(s) to the cell periphery. Middle and right: immunostaining of DC-T cell conjugates against α-tubulin (red) and 3D reconstruction of MTs in DCs (red) and T cells (gold). Nuclei were counterstained with DAPI (blue) and after 3D reconstruction displayed in different shades of blue in DCs (dark blue) and T cells (light blue). (**E-G)** Optimized search and capture of IS dictating optimal centrosome positioning. (**E**) A schematic representation of the simulation model, involving dynamic MTs (blue) emanating from the centrosome (yellowish) and searching for the IS (cyan) located on cell surface. (**E**, *i, ii)* Visual representation and associated probabilistic considerations of the MT dynamics upon hitting the nuclear and cell surface, respectively. The angle *α* determines the chances of MTs dissociation from nuclear surface with increased MT curvature along the nuclear surface. The parameter, *β*, dictates the chances of cell surface-induced instant catastrophe of MTs. (**E**, *iii*) Modulation of the catastrophe frequency of the MTs gliding along the cell surface. (**F**) Average search time < *T_search_* > vs *d_CS−NS_* for a centrally located nucleaus (*d_offset_* = 0) and for different *λ_l_* that modulates MTs catastrophe along cell surface, compared with the scenario where MTs are not allowed to glide along the cell surface (no cell surface gliding). (**G**) Plot of < *T_search_* > vs *d_CS−NS_* for *d_offset_*= 0, 6 µm, and 12 µm, respectively, and without MTs gliding along the cell surface. The optimal centrosome positions denoted by the red star marks are schematically shown on the right. The horizontal blue dashed lines represent the position of the cell’s mid-plane. (**H**) Optimized values of the centrosome distance from the cell center, *h_CS_*, and from the nuclear surface, *d_CS−NS_*, corresponding to the minimum of average search time, <*T_search_*>, for different off-centered positions of the nucleus (see Fig. 8G).

### Modeling highlights a critical role of MT dynamics for centrosome positioning in DCs

Next, we sought to understand how MT dynamics may affect centrosome positioning in DCs and aimed to determine the optimal centrosome position that minimizes the time required for searching the IS in DCs. To explore this, we adopted the well-established “search and capture” hypothesis, initially proposed in the context of mitotic spindle assembly, which leads to the binding of MT filaments to the kinetochores of sister chromatids (Heald & Khodjakov, 2015; Hill, 1985; Prosser & Pelletier, 2017). This framework was further extended in a recent study to examine the docking efficiency of MTs at the IS in T cells (Sarkar *et al*, 2019) and involves dynamically unstable MTs originating from the centrosome, exploring the three-dimensional cellular volume to locate and capture the target IS. We accommodate this established model for MT dynamics in T cells with morphological differences of the MT network of T cells and DCs: T cells are spherical (radius, *r* ≈ 5 *μm*), much smaller than DCs, with a relatively large nucleus (*r* ≈ 3 *μm*), such that the centrosome is generally located close to cell surface and MTs emanating tangentially to the surface (Yi *et al*, 2013; Horňák *et al*, 2020). DCs are large cells (*r* ≈ 18 *μm*) with a relatively smaller nucleus (*r* ≈ 6 *μm*) and centrosome(s) that are typically positioned close to the cell center. This specific arrangement allows MTs to emanate astrally towards the surface, rarely touching it tangentially as in T cells (**Fig. 8D**). Consequently, we incorporate a reduced MT gliding probability into our model: When growing MTs hit the cell surface, they can undergo instant catastrophe determined by the angle of contact with the surface or glide along the surface with catastrophe frequency increasing with gliding distance, as depicted in **Fig. 8E (ii), (iii)**. Note that such cortex-induced changes in MT growth direction were reported in animal cells (Oakley & Brunette, 1993; Waterman-Storer & Salmon, 1997), plant cells (Dixit & Cyr, 2004), yeast (Foethke *et al*, 2009), and HeLa cells (Picone *et al*, 2010) and are well established in theoretical modeling (Sarkar *et al*, 2019; Mallick *et al*, 2022). We further hypothesized that MT growth can be similarly directed by the nuclear surface since we frequently observe MTs in the “geometric shadow” behind the nucleus (the region not accessible to straight MTs growing from the centrosome; see also Fig. 3B; right) and incorporated it into our model as sketched in **Fig. 8E (i)**: When MTs reach the nuclear surface, they first glide along it and subsequently detach at a rate that increases with MT curvature along the nuclear surface. After detachment, MTs grow tangentially from the nuclear surface toward the cell surface seeking the target. To explore how the MT’s gliding ability along the cell surface affects centrosome positioning in DCs, we plot the average search time (< *T_search_* >) against the distance of the centrosome from the surface of a centrally located nucleus, *d_CS−NS_*, while varying the parameter *λ_l_* (**Fig. 8F**) regulating the sensitivity of the catastrophe frequency with MT’s gliding distance (see **Fig. 8E, iii**). A higher *λ_l_*corresponds to a faster MT catastrophe and *vice versa*. We find that for higher *λ_l_*, the average search time is minimized when the centrosome is located near the nuclear periphery. This observation correlates with the scenario where MTs are not allowed to glide along the cell surface, resulting in instant catastrophe upon hitting the cell surface. However, for very small *λ_l_* values, the optimal centrosome position shifts adjacent to the cell surface. With restricted MT gliding along the cell surface, the search progresses via two pathways: MTs can directly reach the target from the centrosome, and MTs that encounter the nucleus glide along its surface and reach the target on the cell surface after detaching from the nuclear surface. In this case, MTs can capture the target most efficiently when they are close to the nuclear surface, minimizing the average distance between the centrosome and the target IS. For centrosome(s) far off from the nucleus, capturing IS located on the cell surface well below the cell’s equatorial plane (hidden below the nucleus) relies on the MTs originating from the centrosome hitting the nuclear surface and gliding along it before reaching the target IS. This in turn increases the overall search time for centrosomes distant from the nucleus. Conversely, if MTs can glide along the cell surface (small *λ_l_*), placing the centrosome near the cell surface accelerates the target capture. Note that, given the experimentally consistent optimal centrosome positioning without MT gliding along the cell surface (or with larger *λ_l_*), henceforth, we proceeded with simulations, preventing the MTs from gliding along the cell surface.

Next, we explored the sensitivity of our findings to off-centered nuclear positioning (**Fig. 8G**). In line with geometric predictions, our results indicate the optimal perinuclear position of the centrosome for a centrally located nucleus and in the vicinity of the cell center for an off-centered nucleus (**Fig. 8G,H**). Surprisingly, even with severely shorter MTs (∼ 30% of the average MT length considered in the model), the optimal centrosome position remains consistent, with prolonged search times (**Fig. EV5F-H**). Evidently, for a largely off-centered nucleus (*d_offset_* = 12 µm), a perinuclear centrosome results in a significant increase in the distance to targets on the cell surface away from the nucleus, leading to extended search times. In contrast, MT arrays from a centrally located centrosome efficiently capture targets across the cell surface due to the minimum average distance to the centrosome. Interestingly, like spherical cells, flattened DCs display optimal centrosome positioning at the cell center for an off-centered nucleus and in the perinuclear region along the short axis of the cell for a centrally located nucleus (**Fig. EV5I-K**).

### Clustering of multiple centrosomes enhances MT search efficiency

To further investigate how centrosome clustering affects the search process, we performed simulations involving four centrosomes, with an evenly distributed array of MTs (**Fig. 9, A-C**). These centrosomes were separated and randomly positioned around the optimal centrosome position determined earlier for a centrally located and off-centered nucleus (see **Fig. 8G,H**). Strikingly, our findings revealed that irrespective of the nuclear positioning, the average search time is minimized when all the centrosomes are clustered together. Clustered centrosomes promote efficient capture of the IS by a unified radial array of MTs. The collective search conducted by MTs nucleating from all the centrosomes originating from a single optimal position increases the likelihood of capturing the target by MTs from at least one of the centrosomes. However, in the case of de-clustered centrosomes, the capture process is not as efficient since the MTs do not originate from the same optimal position. Fewer MTs from each of the centrosomes searching independently for distant targets on the cell surface decreases the chance of a successful capture which effectively increases the overall search time.

**Figure 9.**
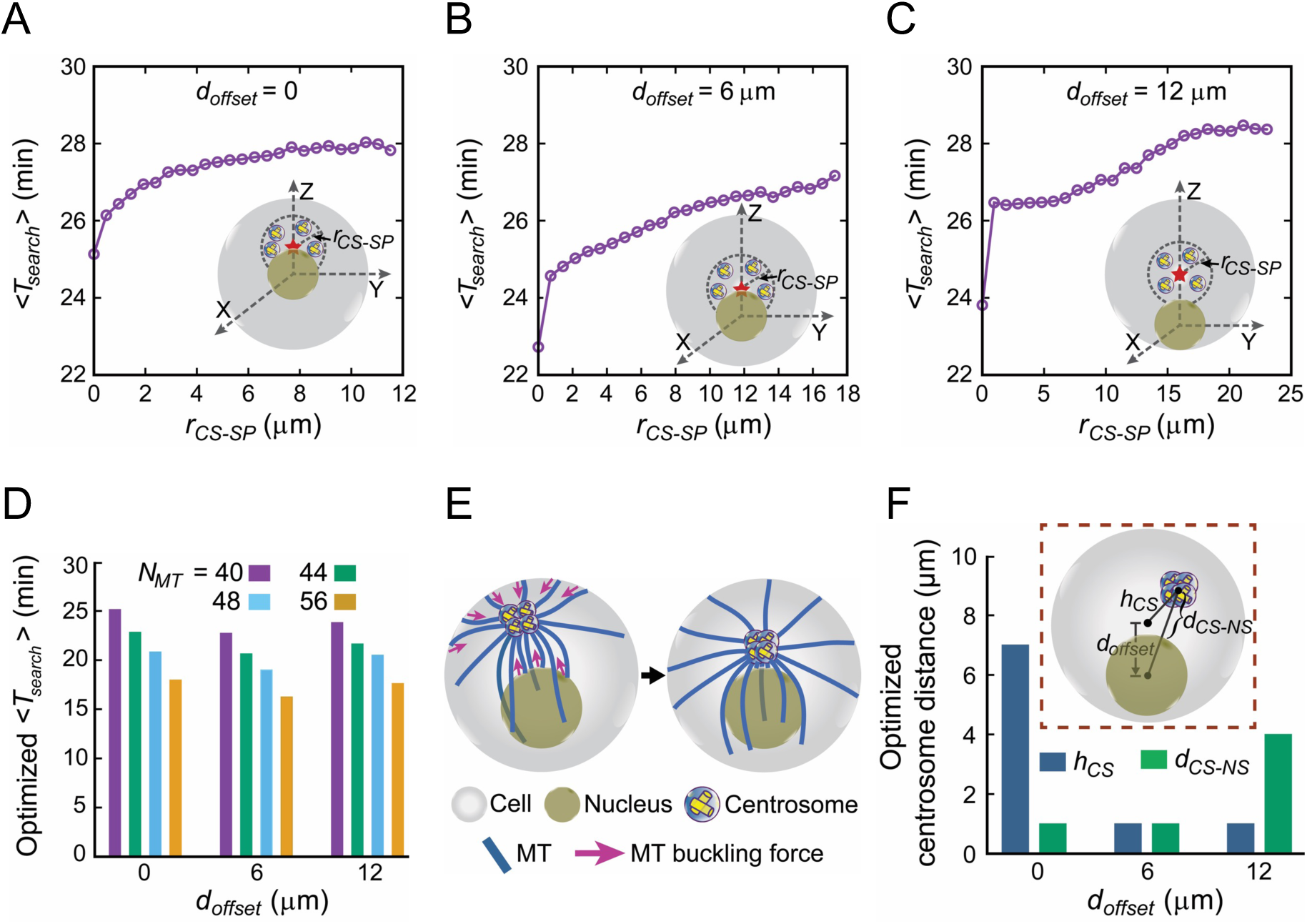
Modeling T cell priming in the presence of multiple centrosomes. (**A-C**) Clustering of supernumerary centrosomes promotes optimized search. Average search time < *T_search_* >, is plotted against *r_CS−SP_* for different off-centered positions of the nucleus and without MTs gliding along the cell surface. *r_CS−SP_* is the radius of the imaginary sphere centered at the optimal centrosome position (denoted by red star marks) obtained in **Fig. 8G,H** within which centrosomes are randomly placed. The insets provide illustrations of nuclear positioning and the placement of centrosomes around optimal positions for various off-centered positions of the nucleus. (**D**) Average search time < *T_search_* > vs *d_offset_* with all the centrosomes clustered together (*r_CS−SP_* = 0) and for different number of centrosomal MTs. (**E-F**) A mechanistic force balance model, considering MT’s interaction with the nuclear and cell surfaces, supports observed centrosome positioning in DCs. (**E**) A schematic representation of the positioning of four closely placed centrosomes in DCs, governed by pushing forces generated by MT buckling at the cell and nuclear surface. (**F**) The final equilibrium distance of the centrosome-cluster from the nuclear surface, *d_CS−NS_* and cell center, *h_CS_* (see the inset for a schematic depiction) for *d_offset_* = 0, 6 µm, *d_offset_* = 0, 6 12 µm, respectively.

Noting that our experiments revealed a ∼10 − 20% increase in the total number of MTs in DCs containing multiple clustered centrosomes compared to DCs with a single centrosome, we further employ our computational model to investigate the correlation between MT search time and the number of MTs. Interestingly, we find a decrease in the search time with an increasing number of MTs (**Fig. 9D**). This behavior is consistent with our analytical prediction of average search time without a nucleus, estimated within the model framework introduced earlier (Sarkar *et al*, 2019) (see *Appendix*). Note that, dispersing the centrosomes may also facilitate faster target capture when MTs are allowed to glide freely along the cell surface rather than restricting their growth (**Fig. EV6A-C**). Under such conditions, the dispersed centrosomes allow MTs to efficiently capture the targets by gliding along the cell surface, resulting in a reduced search time. Overall, our model highlights the pivotal role of multiple clustered centrosomes, coupled with increased MT numbers, efficiently activating T cells by facilitating quicker access to the IS through dynamic MTs.

### A mechanistic force-balance model exhibiting centrosome positioning in DCs

To find the mechanically stable configuration of multiple centrosomes, irrespective of a putative attractive force between them, we used an established mechanistic model of MT-generated forces resulting from interactions with the nucleus and the cell surface (Letort *et al*, 2016; Som *et al*, 2019; Zhu *et al*, 2010). While pushing against the cell or nuclear surfaces, a polymerizing MT is buckled, and the corresponding buckling force is applied on the centrosomes. The resultant force causes the centrosomes to move through the cytoplasm (Zhu *et al*, 2010; Som & Paul, 2023; Som *et al*, 2019) as sketched in **Fig. 9E**. Similar to the search and capture model described earlier with frequent observations of MTs at the “geometrical shadow” behind the nucleus (see also Fig. 3B, right), we hypothesize that MT growth can be directed by the nuclear surface. Accordingly, our model assumes that the majority of MTs slide along the nuclear surface and only a small fraction (∼ 10%) buckles at the nuclear surface, whereas all MTs hitting the cell surface generate buckling-induced-pushing force on the centrosomes.

We observe that the centrosome-cluster stayed together and positioned along the line joining the nuclear center and the cell center. The clustered centrosomes remain close to the nuclear surface (∼1 μm away from the nuclear surface) when the nucleus is fixed at the cell center (*d_offset_* = 0) and near the cell center when the nucleus is off-centered (*d_offset_* = 6 µm and 12 µm) (**Fig. 9F** and **movies EV5-7**). These findings qualitatively corroborate our experimental observations and simulation results of the optimal centrosome positions estimated geometrically and by the search-and-capture scheme. As expected, when more MTs are assumed to buckle at the nuclear surface, the distance between the centrosome cluster and the nuclear surface increases due to the higher pushing forces generated on the centrosomes from the nuclear surface (**Fig. EV6D**).

Altogether, our computational approach highlights a beneficial role for multiple clustered centrosomes in APCs, which nucleate a larger number of MT filaments and collectively stay close to the cell center to optimally form cell-cell contacts and ultimately activate T cells.

## Discussion

Mounting of an efficient immune response requires specific cell-cell interactions, which take place in specialized immune compartments such as secondary lymphoid organs. During the past decades much progress has been made in elucidating the dynamic behavior of cell surface and intracellular signaling molecules associated with IS formation and T cell activation, while much less is known about the dynamic changes within APCs during TCR engagement.

DCs are critical players of the innate immune system, which activate T cells via antigen presentation. Upon antigen encounter, a proportion of DCs undergoes an incomplete mitosis leading to the accumulation of extra centrosomes and doubling of DNA content during the subsequent G1 phase (Weier *et al*, 2022). In addition, overduplication of centrioles results in the presence of diploid cells carrying extra centrosomes. DCs possessing amplified centrosomes exhibit elevated levels of cytokine secretion, an increased capacity to activate T cells as well as enhanced persistent locomotion in response to chemotactic gradients (Weier *et al*, 2022). Of note, enhanced effector functions of immune cells in which centrosomes have been artificially amplified by PLK4 overexpression, were also observed in B cells processing and presenting antigens and in microglia phagocytosing dead neurons (Yuseff *et al*, 2011; Möller *et al*, 2022). Overall, these findings imply, that amplification of centrosomes may contribute to regular cell and tissue physiology within the immune compartment to enhance specific effector functions in a context-dependent manner and pose the question whether specific cell-cell interactions are modulated by the presence of extra centrosomes.

Here we demonstrate on a single cell resolution that DC centrosome and MT integrity are key for efficient induction of T cell proliferation, while centrosome repositioning in DCs to the IS is dispensable for CD4^+^ T cell activation. Moreover, we provide evidence that amplified centrosomes enhance T cell activation capacity of DCs by clustering extra centrosomes close to the cell center and the nucleus. In line with these findings, similar observations were described, which analysed APC-CD4^+^ T cell synapses in the absence of DC MTOC polarization to the synapse (Bouma *et al*, 2011; Mittelbrunn *et al*, 2009). These studies further focused on the DC actin cytoskeleton which is reorganized during DC-T cell contacts and controls contact duration and priming efficiency of T cells (Leithner *et al*, 2021). In parallel to T helper synapses, other studies have suggested that T cell centrosome integrity is required for efficient cytotoxic T cell-mediated killing (Tamzalit *et al*, 2020). Intriguingly, it is controversially discussed whether T cell centrosome polarization is dispensable or required for trafficking and directional secretion of lytic granules towards the cytotoxic synapse (Bertrand *et al*, 2013; Tamzalit *et al*, 2020; Chauveau *et al*, 2010). One reason why intracellular MTOC reorganization upon cell-cell contact formation differs in immune cells might be that smaller cells such as T- and B cells need to reorient their centrosome due to geometric hindrance of the nucleus in order to efficiently deliver cargos via MT filaments to the contact zone. By contrast, larger cells such as DCs exhibit a centrally localized centrosome, which forms astral microtubule arrays that reach out to the cell membrane without being strongly disturbed by the presence of a comparably small nucleus. Moreover, DCs form multi-conjugated synapses which make a centrally localized centrosome advantageous for minimizing the average MT search time to reach every point on the cell membrane that are potential sites to form additional cell-cell contacts. Consequently, centrosome re-orientation to one side would increase MT search time to form contacts at opposite sites. In this context, the connection between MT dynamics and centrosome positioning in DCs was further established by our computational model.

Promoting gliding of MTs along the nuclear surface while restricting it along the cell surface we demonstrate a consistent and optimal centrosome position close to the cell center for off-centered positioning of the nucleus and right above the nuclear surface for a centrally located nucleus. Our model construction is in tune with the observed MT arrangement in DCs, presenting a limited number of MTs growing along the cell surface, and a significant number of them surrounding the nuclear surface. Whether the hindrance of MT growth along the cell surface is attributed to the surface ruggedness of DCs or whether it involves the potential role of MT-end binding proteins actively promoting MT catastrophe along the surface (Foethke *et al*, 2009; Varga *et al*, 2009; Wu *et al*, 2006), requires further experiments. Interestingly, enhanced MT sliding along the cell surface yields a very different outcome. It changes the optimal centrosome position from perinuclear to near the cell surface. The outcome is significant in MT-driven search processes, where stable MTs are guided by the topology of the cell surface. For instance, in T cells, which share the common IS with DCs, MTs appear to predominantly slide and curve along the cell surface while approaching the IS. The dynein molecules residing at the IS capture the approaching MT filaments and facilitate centrosome relocation toward the IS by generating tension on the MT (Combs *et al*, 2006; Kuhn & Poenie, 2002; Yi *et al*, 2013). Our findings underscore the adaptability of immune cells regulating their internal organization to generate immune responses specific to particular cell types.

DCs carrying multiple centrosomes demonstrated a propensity for tight clustering of centrosomes during antigen-specific DC-T cell interactions. Our computational study, in line with experiemental observations, further illuminated that during the activation of multiple T cells, clustering of centrosomes in DCs led to efficient capture of the IS by dynamic MTs, accelerating the activation of T cells. Previous studies highlighted the role of a linker composed of Rootletin and C-Nap1 in centrosome tethering, physically holding the centrosomes together during interphase and dissociating during the onset of mitosis, enabling centrosome separation (Bahe *et al*, 2005; Faragher & Fry, 2003; Fry *et al*, 1998; Mardin *et al*, 2010). A recent study pointed to the role of the kinesin-14 motor protein Kif25 in coalescing supernumerary centrosomes into a single pole (Decarreau *et al*, 2017). Intriguingly, our study unveiled a novel mechanism for positioning tightly clustered multiple centrosomes during interphase in DCs, involving the dynamic interaction of MTs with the cell and nuclear surfaces. Our findings reveal that due to these mechanical forces, centrosomes remain clustered throughout their temporal evolution, without requiring any explicit molecular interactions between the centrosomes. Multiple pathways likely coordinate to lay down a robust clustering of the centrosome crucial for the functioning of DCs.

From a clinical perspective, centrosomal clustering is emerging as a novel strategy to specifically target cancer cells, which frequently harbor extra centrosomes referred to as centrosome amplification (Sato *et al*, 1999; Pihan *et al*, 1998, 2003; Krämer *et al*, 2005). To avoid spindle multipolarity, cancer cells cluster amplified centrosomes during cell divisions in order to form a pseudo-bipolar spindle configuration (Ganem *et al*, 2009; Chatterjee *et al*, 2020). However, transient centrosome de-clustering leads to the formation of a multipolar spindle intermediate and mis-segregation of chromosomes (Ganem *et al*, 2009). In this context, de-clustering agents are currently tested in pre-clinical and clinical trials as centrosome de-clustering induces multipolar spindle formation and subsequent cell death (Castiel *et al*, 2011; Firdous *et al*, 2023). The complete mechanism(s) how centrosome clustering is regulated in cancer cells is still unresolved and requires further investigations.

Overall, our study emphasizes the pivotal role of (extra) centrosomes and the MT cytoskeleton for eliciting efficient immune responses. Moreover, it highlights that amplified centrosomes are not necessarily a threat under homeostatic conditions and challenges the current view of extra centrosomes to be exclusively present in transformed cancer cells. Our results will promote future work on amplified centrosomes and their role in immune cells for performing fundamental effector functions.

## Materials and Methods

### Mice

All mice used in this study were bred on a C57BL/6J background and maintained at the institutional animal facility in accordance with the German law for animal experimentation. Permission of all experimental procedures involving animals was granted and approved by the local authorities (Landesamt für Natur, Umwelt und Verbraucherschutz North Rhine-Westphalia [LANUV NRW under AZ81-02.05.40.19.022]). CETN2-GFP and Nur77^GFP^ mice were purchased from Jackson (CB6-Tg(CAG-EGFP/CETN2)3-4Jgg/J and B6N.B6-Tg(Nr4a1-EGFP/cre)820Khog/J). OVA-specific OT-II mice were a gift of Sven Burgdorf.

### Dendritic cell culture

Femurs and tibias from legs of 3–5 month-old CETN2-GFP expressing mice were removed and placed in 70% ethanol for 2 min. Bone marrow was flushed with PBS using a 26 gauge needle. 2*10^6^ cells were seeded per 100 mm petri dish containing 9 ml of complete medium (Roswell Park Memorial Institute (RPMI) 1640 supplemented with 10% Fetal Calf Serum, 2 mM L-Glutamine, 100 U/ml Penicillin, 100 μg/ml Streptomycin, 50 μM ß-Mercaptoethanol; all Gibco) and 1 ml of Granulocyte-Monocyte Colony Stimulating Factor (GM-CSF, supernatant from hybridoma culture). On day 3 and 6 complete medium supplemented with 20% GM-CSF was added to each dish. To induce maturation, cells were stimulated overnight with 200 ng/ml lipopolysaccharide (LPS) and used for experiments on day 8-9. Alternatively, DCs were frozen in FCS containing 10% DMSO on day 7. For experimental use they were thawed the day before the experiment and stimulated with 200 ng/ml LPS overnight.

To prevent new pro-centriole formation cells were cultured in the presence of the PLK4 inhibitor Centrinone (Tocris; 250 nM or 500 nM) or control (solvent DMSO) during differentiation and maturation. To induce microtubule depolymerization, cells were treated with 1 µM pretubulysin for 1 h after antigen-loading. Cells were washed (wash-out) or not (w/o wash-out) with full media before T cell addition. To induce centrosome de-clustering PJ-34 (Sigma-Aldrich) was used. Cells were loaded with OVAp for 2 h and subsequently treated with 50, 100 or 200 µM of PJ-34 or control (solvent H_2_O) for 3 h. Cells were washed two times with full media and incubated with OT-II-specific T cells at the indicated time points.

### Flow cytometry

For flow cytometric analysis, cells were washed with PBS once and incubated 10 min with anti-CD16/CD32 antibody (1:100) in blocking buffer (1x PBS, 1% BSA, 2 mM EDTA). Staining with fluorescently labelled antibodies diluted in blocking buffer was carried out for 20 minutes at 4°C. The following antibodies were used: hamster anti-mouse CD11c-PE (N418, 1:500), rat anti-mouse MHCII (I-A/I-E)-APC-Cy7 (M5/114.15.2, 1:800), rat anti-mouse CD4-APC (RM4-5, 1:500), rat anti-mouse CD19-PE (6D5, 1:500), rat anti-mouse MHCII (I-A/I-E)-PE-Dazzle (M5/114.15.2, 1:600), hamster anti-mouse CD69-FITC (H1.2F3, 1:200), hamster anti-mouse CD69-PE-Dazzle (H1.2F3, 1:200), rat anti-mouse CD62L-PE-Cy7 (MEL-14, 1:500). For live dead staining DR (1:500) was used. Afterwards, cells were washed once with blocking buffer and data acquired at the LSRII flow cytometer (BD Bioscience). Data analysis was performed using FlowJo v10.8.1.

To determine T cell activation via CD69 upregulation and CD62L downregulation, DC co-cultures with splenocytes were analysed after 20-22 hours via flow cytometry as described above. T cell proliferation rates were assessed by dilution of CFSE. Therefore, prior to incubation with DCs, splenocytes were stained with a final concentration of 0.5 µM CFSE (Invitrogen) for 7 minutes at 37°C in PBS and washed with complete medium. Co-cultures were incubated for 2.5 days. Nur77-dependent GFP upregulation was also determined via flow cytometry after the indicated timepoints of naïve CD4^+^ T cell co-culture with DCs.

### Sorting of 2N BMDCs

To sort DCs based on their DNA content, mature BMDCs were harvested and stained with Vybrant DyeCycle Violet Stain (1:1000, Thermofisher; V35003) in RPMI without phenol red for 20 min at 37°C. Cells were sorted at the ARIAIII Sorter (BD Bioscience) according to their ploidy level with focusing on diploid cells (2N) and dismissing polyploid cells. Cells were re-analysed after the sort to ensure purity of the individual subpopulations. Afterwards, cells were recovered in full medium at 37°C for at least 30 minutes.

### Mixed lymphocyte reactions (MLR)

For DC-T cell co-culturing assays sorted DCs (1-2*10^4^ cells/well) were seeded in 96-well U-bottom plates. After recovering time of 30 minutes, cells were incubated with OVAp antigen (OVA_323-339_: specifically recognized by CD4^+^ OT-II T cells; 0.01 µg/mL, 0.1 µg/mL, 1 µg/mL) or without OVAp (controls) for 2 hours. In the meantime, splenocytes were isolated from OT-II mice or Nur77^GFP^/OT-II mice. Therefore, splenic, inguinal, axillary and brachial lymph nodes were removed and smashed through a 70 µm filter using PBS and a syringe piston. After centrifugation (400 x g 5 minutes), ACK lysis buffer (Gibco, A10492-01) was added for 5 minutes at RT before stopping the red blood cell lysis with PBS containing 2% FCS and 2 mM EDTA. Cells were filtered through a 40 µm filter, centrifuged and adjusted to the respective cell number to co-culture with DCs or for subsequent naïve CD4^+^ T cell isolation. Naïve CD4^+^ T cell isolation was performed according to the manual of the EasySep Mouse Naïve CD4^+^ T Cell Isolation Kit (STEMCELL, 19765). After removing the antigen from the DCs, T cells were added to DCs in a ratio of 1:2 or 1:5.

### Under-agarose interaction assay

To allow visualization of centrioles during live cell imaging DC-T cell co-cultures were injected under a block of agarose as previously described (Leithner *et al*, 2021). Briefly, a custom-made chamber was built by gluing a 1-cm plastic ring with paraffin into a glass-bottom dish. 1% agarose solution was prepared by mixing 0.2 g UltraPure agarose (Invitrogen, 16500-100) with 5 mL water, 5 mL 2x Hanks’ Balanced Salt Solution (HBSS 10x, Gibco, 14185052) and 10 mL phenol red-free RPMI1640 Medium supplemented with 20% FCS and 1% penicillin 100 U/mL/streptomycin 100 µg/mL. For live cell imaging, ascorbic acid was added to a final concentration of 50 µM. 500 µL of the heated agarose was poured into each chamber. After polymerization, the dishes were filled with water around the agarose and incubated for 45-60 minutes at 37°C, 5% CO_2_ to equilibrate the agarose. For the experiment, cells were injected with a small tip under the agarose in a volume of 0.4-0.6 µL. To allow visualization of interaction during live cell imaging T cells were stained with the Calcium-sensitive dye Cal520 (Abcam) prior to injection. For efficient staining, cells were incubated with 3 µM Cal520 in phenol red-free RPMI1640 medium supplemented with 20% FCS for 30 minutes at 37°C. To avoid toxic effects the dye was efficiently removed by washing two times. Live cell imaging was started directly after DC and T cell injection. Alternatively, to allow immunofluorescence staining, cells were fixed after 60-90 min in the incubator with 4% paraformaldehyde (PFA) solution overnight at 4°C. The next day, agarose was removed carefully and the cells were washed with PBS two times before staining.

### Immunofluorescence

Cells were immobilized by incubating 1-2 µL of cell suspension on uncoated cover slips for 5 minutes at 37°C before adding 4% PFA for 20 minutes. Fixed cells were washed twice with PBS for 10 minutes. To allow intracellular antibody staining cells were permeabilized adding 0.2% Triton X-100 (Sigma) in PBS for 30 minutes at room temperature. After washing 3 x 10 minutes with PBS samples were incubated in blocking solution (5% BSA (Roth) in PBS) for 1 hour to prevent unspecific binding of the antibodies. Next, samples were incubated with primary antibodies diluted in blocking solution over night at 4°C. Afterwards, cover slips were washed 3 x 10 minutes with PBS and stained with secondary antibodies diluted in blocking solution in the dark for 1 hour at room temperature. After three times washing 10 minutes with PBS cover slips were mounted with DAPI-containing mounting medium (Invitrogen) and sealed with nail polish before imaging.

The following primary antibodies were used: rat anti-mouse alpha-tubulin (YL1/2, Invitrogen, 1:500), mouse anti-mouse acetylated-tubulin (6-11B-1, Sigma-Aldrich), mouse anti-mouse γ-tubulin (GTU-88, Sigma-Aldrich, 1:500), rabbit anti-mouse γ-tubulin (polyclonal, Abcam, 1:500), rabbit anti-mouse CDK5RAP2 (polyclonal, Sigma-Aldrich, 1:500).

The following secondary antibodies were used in a dilution of 1:400: Donkey Anti-Mouse Alexa Fluor 647 AffiniPure F(ab’)_2_ Fragment IgG (H+L), Donkey Anti-Mouse Cy3 AffiniPure F(ab’)_2_ Fragment IgG (H+L), Donkey Anti-Rat Cy3 AffiniPure F(ab’)_2_ Fragment IgG (H+L), Donkey Anti-Rabbit Alexa Fluor 647 AffiniPure F(ab’)_2_ Fragment IgG (H+L), Goat Anti-Rabbit Cy3 AffiniPure F(ab’)_2_ Fragment IgG (H+L) (all from Jackson ImmunoResearch).

### Microscopy

Confocal microscopy was performed on a motorized stage at RT with an inverted microscope equipped with an Airyscan module; a Plan-Apochromat 63×/1.4 oil DIC objective; 488, 561, and 633 laser lines; and a photomultiplier tube (all Zeiss). For fixed samples 0.2 µm sections were acquired leading to Z-stacks of mostly 4-8 µm height. To analyse MT filaments, images were acquired using the Airy module and posttreated by deconvolution. The same confocal imaging set up was used for live cell imaging. During live-cell acquisition dishes were placed in a 37°C chamber and cells imaged at a 10 to 20 seconds interval for 30-60 minutes. The auto-focus option was used to keep the centrioles in focus. For all experiments, imaging software ZEN Black 2.3 SP1 was deployed.

#### Image analysis

Image processing and data analysis were performed using ImageJ. For measurements of distances between or from centrioles, centrioles were tracked by using the Manual Tracking plugin. Quantitative intensity measurements were carried out measuring RAW integrated density or integrated density of selected regions of interest. For determining MT numbers emanating from the centrosome, MT filaments were counted manually at a defined area around the centrosome. The whole image *z-*stack was used to precisely identify individual MT filaments. 3D reconstruction of MT filaments within DCs and T cells was performed with IMARIS 9 software.

### Expansion microscopy (ExM)

Control or Centrinone-treated BMDCs were seeded on poly-L-Lysin (0.1 mg/ml, Sigma) coated coverslips and allowed to adhere for 1h at 37°C. U-ExM protocol was carried out as described previously (Laporte *et al*, 2022; Gorilak *et al*, 2021) with mild adjustments. Cells were fixed with 4% PFA in PBS at 37°C for 15 minutes, following post-fixation with 0.7% PFA and 2% acrylamide (AA) at RT overnight followed by PBS wash prior to gelation. Gelation was carried out in a wet chamber placed on ice and covered with Parafilm. Washed coverslips were transferred promptly on a small drop of gelation solution (19% sodium acrylate, 10% AA, and 0.1% N, N’-methylenebisacrylamide initiated with 0.5% tetramethylethylendiamine and 0.5% ammonium persulfate). Gelation was initiated on ice for 5 minutes and then transferred to 37°C for 30 minutes to allow polymerization. Gels were then detached from the coverslips in a denaturation buffer (50 mM Tris-base, 200 mM NaCl, 200 mM SDS in ddH_2_0, pH 9) and denatured at 95°C for 1 hour. Gels were expanded with 3x ddH_2_O wash and a small piece of gel was cut out for staining. Staining was performed sequentially overnight at RT in a staining buffer (2% BSA, 10% sodium azide in PBS) with 3x ddH_2_0 wash between each step with the following primary: mouse anti-mouse acetylated-tubulin (C3B9, 1:10), rabbit anti-mouse γ-tubulin (polyclonal, Abcam, 1:200), rabbit anti-mouse CDK5RAP2 (polyclonal, Sigma-Aldrich, 1:200) and secondary: goat anti-rabbit Alexa Fluor 488 IgG H+L (1:500, A11008) and goat anti-mouse Alexa Fluor 555 IgG H+L (1:500, A21422) antibodies. NHS Ester Atto 425 (20 mg ml^−1^ in PBS, ATTO-TEC) staining was done in PBS at RT for 1.5 hours and gels were expanded with 3x ddH_2_0 wash before imaging. Microscopy was performed on a Nikon Eclipse Ti2 microscope equipped with Yokogawa CSU-W1 spinning disc module using the CF Plan-Apochromat VC 60×/1.2 water objective.

### Statistical Analysis

Data analysis was carried out with GraphPad Prism 10 (GraphPad Software, San Diego, CA, USA). Samples were tested for Gaussian distribution using D’Agostino-Pearson omnibus normality test to fulfil the criteria for performing Student’s *t*-tests. Welch’s correction was applied when two samples had unequal variances. When data distribution was not normal, Mann-Whitney test was carried out. For small data sets, Gaussian distribution was assumed but could not be formally tested. For analysis of Nur77^GFP^ expression, CETN2-GFP^low^ and CETN2-GFP^high^ samples from individual experiments were paired. For multiple comparisons where data distribution was normal, one-way ANOVA was used followed by Dunnett’s multiple comparisons as post-hoc test. When data distribution was not normal, Kruskal-Wallis test with Dunn’s multiple comparisons was used. All graphs display mean values ± s.d. (95% Confidence Interval). No statistical method was used to predetermine sample size. The experiments were not randomized and investigators were not blinded to allocation during experiments and outcome assessment. Individual experiments were validated separately and only pooled if showing the same trend. The level of significance was denoted as *, *P* < 0.0332; **, *P* < 0.002; ***, *P* < 0.0001 and ****, *P* < 0.00001 as indicated in the figure legends.

### Computational model

DCs are modeled as spheres with radius *R_cell_* containing a spherical nucleus of radius *R_nu_* as depicted in Fig. 8A. The line connecting the nucleus center and cell center defines the z-axis. The centrosome is placed along the z-axis at a distance *h_CS_*, above the cell center and *d_CS−NS_* (= *h_CS_* − *R_nu_*) away from the nuclear surface. The parameter *d_offset_* determines the distance of the nucleus center from the cell center as depicted in Fig. 8C.

### Average geometrical distance between centrosomes and the target points on cell surface

The optimal geometric centrosome position is determined by minimizing the average distance, < *d_short_* >, between the centrosome and all target points on the cell surface. In the absence of a nucleus, this distance is the shortest straight geometric distance between the centrosome and the target points on the cell surface. In the presence of a nucleus, a region of the cell surface lies in the “geometric shadow” behind the nucleus, which is the region not accessible to straight MTs growing from the centrsosome. In such configurations, two tangents are drawn on the nuclear surface, one originating from the centrosome and the other from the target point on the cell surface, ensuring that the corresponding tangent points on the nuclear surface, centrosome, and the centers of the nucleus and target remain on the same plane. The total distance is then determined as the sum of the lengths of the tangential segments and the intermediate arc length along the nuclear surface (see schematic in Fig. 8A). The average geometric distance, < *d_short_* >, is then computed by averaging over 10^6^ random target points on the cell surface.

### Average search time

The optimal dynamical centrosome position is determined by the minimum of the search time, *T_search_*, required for dynamic MTs to capture a target located randomly on the cell surface. The target zone is represented as a circular disk of radius *R_τ_* embedded in the cell surface, as illustrated in Fig. 8E. The target is located on the cell surface at an arbitrary position, specified by the polar angle *θ_τ_* (∈ 0 − 180^3^) and azimuthal angle *φ_τ_* (∈ 0 − 360^3^), measured from the positive z-axis and x-axis, respectively. A specific number of *N_MT_* MTs nucleate from the centrosome and explore the surrounding three-dimensional cellular space searching for the target. In the presence of supernumerary centrosomes, the total *N_MT_* MTs are distributed evenly among all the centrosomes. The dynamically unstable MTs exhibit consistent growth at a velocity, *v_g_*, switch to a shrinking phase with a catastrophe frequency, *v_s_*, and shortening at a different velocity, *f_r_*. The catastrophe frequency is chosen such that, on average, MTs could cover a distance equivalent to half of the cell perimeter. The simulations are performed with zero rescue frequency (*f_r_*), preventing shrinking MTs from switching to the growth phase. A zero rescue frequency is optimal since it minimizes the search time that MT spends exploring directions lacking the target (Holy & Leibler, 1994; Sarkar *et al*, 2019; Wollman *et al*, 2005; Paul *et al*, 2009). A Monte Carlo algorithm is implemented to simulate individual MTs. At each computational time step (Δt), a uniform random number between 0 and 1 was compared with the probability 1 − *exp*(−*f_c_* Δ*t*) that an MT switches from growth to shortening. If the random number is found to be less than this probability, the MT begins shortening. Once an MT shrinks back to the centrosome, a new growth cycle starts in another random direction.

The direction of MT nucleation from the centrosome was governed by two angles, polar angle, *θ* (∈ 0 − 180^3^) and azimuthal angle, *φ_τ_* (∈ 0 − 360^3^), in the standard spherical polar coordinate system. Depending on the direction of nucleation and subsequent interactions with the cell or nuclear surface, two distinct scenarios arise:

i. The model assumes that when MTs interact with the nuclear surface, they can move along the nuclear surface maintaining the same azimuthal angle *φ* at which they originated from the centrosome. During this movement along the nuclear surface, the increasing curvature of the MTs can promote their detachment at a rate 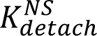. After detachment, the MTs grow tangentially from the nuclear surface and approach the cell surface searching for the target. The rate of detachment, 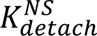, is assumed to be an exponentially increasing function of the MTs’ curvature along the nuclear surface, as depicted in Fig. 8E, (i). Two parameters, *k*_0_ (the prefactor of the exponential term) and *α_d_* (a phenomenological constant within the exponential term) regulate the sensitivity of the MTs’ detachment rate (see Fig. 8E, (i)). Smaller values of *k*_0_ and/or larger values of *α_d_* promotes the MTs to glide along the nuclear surface, minimizing the chances of rapid detachment. MTs that extend beyond half of the nuclear perimeter from their initial contact points are immediately detached from the nuclear surface.
ii. If the MTs reach the cell surface either after detaching from the nuclear surface or directly from the centrosome, they can eventually capture the target if the MT tip hits the target. Otherwise, they move along the cell surface seeking the target or experience catastrophe induced by the surface curvature. This cell surface induced MT catastrophe is incorporated in the model based on previous studies demonstrating that the distribution of MTs along the cell surface is influenced by the cell shape, and ruggedness of the cell surface regulating the MT bending and catastrophe (Laan *et al*, 2012; Picone *et al*, 2010; Pavin & Tolić-Nørrelykke, 2014). The model assumed a probabilistic catastrophe of the MTs at the cell surface depending on the angle of incidence (*β*) of the MT with respect to the local normal. The probability is chosen such that the tangential incidence (*β* = π/2) of MTs would promote their gliding along the cell surface and normal incidence (*β* = 0) would induce catastrophe at a certain rate (Fig. 8E, (ii)). Specifically, the catastrophe probability 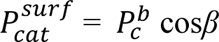 and the gliding probability 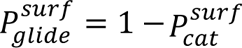 are considered. The value of 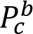 is chosen to be 1 to ensure that MTs experience instant catastrophe for normal incidence on the cell surface (*β* = 0). If the MTs begin moving along the cell surface overcoming the cell surface -induced catastrophe, the MTs’ movement along the cell surface is further restrained by modulating the catastrophe frequency of gliding MTs according to 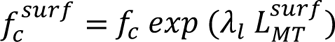, where 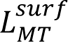 represents the segment of the MT’s length gliding along the cell surface, and *λ_l_* is a phenomenological constant determining the rate at which the catastrophe frequency increased per unit length of the MT (Fig. 8E, (iii)). This additional consideration of catastrophe modulation is motivated by experimental observations of MT organization in DCs, which revealed that a very small fraction of the MTs or almost no MTs appeared to be gliding along the rugged cell surface (Weier *et al*, 2022). The simulation is continued until a successful target-MT attachment is formed. The average search time, < *T_search_* >, is calculated using 10: different random positions of the target on the cell surface. The parameters used in the simulation are tabulated in ***Table S1*** in ***Supplementary Materials***.

### A mechanistic force-balance model demonstrating multi-centrosomal positioning

The centrosomes are considered as small spherical objects with *r_CS_* (∼ 0.5 µm), free to move within the cellular space between the cell surface and a nucleus fixed in space at various positions within the cell. MTs are cylinders of vanishing radii and nucleated from the centrosome uniformly in all directions. The number of MTs per centrosome is 10 (totalling 40 for four centrosomes) and the dynamics of each MT is governed by *v_g_*, *v_s_*, *f_c_* and *f_r_* discussed earlier. Based on earlier studies of centrosome positioning and stability in interphase cells, the model assumes a pushing force-dominated regime arising from MT interaction with the cell and nucleus (Fig. 9E) (Burakov & Nadezhdina, 2013; Som *et al*, 2019; Zhu *et al*, 2010). In our model, a growing MT of length *L_MT_* hitting the cell or nuclear surface buckles as per first-order Euler’s buckling and translates a pushing force 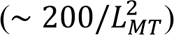 to the corresponding centrosome (Som *et al*, 2019; Zhu *et al*, 2010). The model assumes that only a fraction of MTs hitting the nucleus buckles, while the others continue to slide along the nuclear surface. The buckling force is inversely proportional to the MT length and hence only short MTs generate significant buckling force and also undergo instant catastrophe (Letort *et al*, 2016; Som *et al*, 2019) leading to short-lived MT pushing forces. The resultant total force on the centrosomes can move them through the viscous cytoplasm following Stokes’s law (Som *et al*, 2019; Som & Paul, 2023). If *F_CS_* is the net force acting on a centrosome, *V_CS_* the instantaneous velocity and *μ* the effective viscous drag on centrosome, then as per Stokes’s law *F_CS_* = *μV_CS_* (with *μ* = 6*πηr_CS_*; *η* is the coefficient of cytoplasmic viscosity) (Som & Paul, 2023; Som *et al*, 2019). The position of centrosomes is updated using the coarse-grained time step Δ*t* (= 10^−2^*s*). The parameter values are mentioned in ***Table S1*** in ***Appendix***.

## Acknowledgements

The authors acknowledge the Imaging Methods Core Facility at BIOCEV, institution supported by the MEYS CR (LM2023050 Czech-BioImaging) for their support & assistance in this work. MH and EM thank Vladimir Varga for providing the C3B9 anti-acetylated tubulin antibody. AS and RP acknowledge IACS, Kolkata, for funding and support. EK acknowledges the TRA Life and Health (University of Bonn) as part of the Excellence Strategy of the federal and state governments.

## Author contributions

IS, AW, PK, MH, KS, JB, EM, MHons and EK performed experiments. SU helped with image analysis. UK provided pretubulysin and LS verified its efficiency. HR, AS, SS and RP performed modelling of centrosome configuration and MT dynamics. ZA gave technical support and supervised KS. EK designed and supervised the research. IS, HR, AS, RP and EK wrote the manuscript. All authors discussed the results and implications and commented on the manuscript at all stages.

## Data and materials availability

Data that support the findings of this study are available within the article and its supplementary information or on request from the corresponding author(s).

## References

Bahe S, Stierhof Y-D, Wilkinson CJ, Leiss F & Nigg EA (2005) Rootletin forms centriole-associated filaments and functions in centrosome cohesion. J Cell Biol 171: 27–33

Banchereau J, Briere F, Caux C, Davoust J, Lebecque S, Liu Y-J, Pulendran B & Palucka K (2000) Immunobiology of Dendritic Cells. Annu Rev Immunol 18: 767–811

Benvenuti F, Lagaudrière-Gesbert C, Grandjean I, Jancic C, Hivroz C, Trautmann A, Lantz O & Amigorena S (2004) Dendritic Cell Maturation Controls Adhesion, Synapse Formation, and the Duration of the Interactions with Naive T Lymphocytes. J Immunol 172: 292–301

Bertrand F, Müller S, Roh K-H, Laurent C, Dupré L & Valitutti S (2013) An initial and rapid step of lytic granule secretion precedes microtubule organizing center polarization at the cytotoxic T lymphocyte/target cell synapse. Proc Natl Acad Sci 110: 6073–6078

Bornens M (2012) The Centrosome in Cells and Organisms. Science 335: 422–426

Bouma G, Mendoza-Naranjo A, Blundell MP, Falco E de, Parsley KL, Burns SO & Thrasher AJ (2011) Cytoskeletal remodeling mediated by WASp in dendritic cells is necessary for normal immune synapse formation and T-cell priming. Blood 118: 2492–2501

Braig S, Wiedmann RM, Liebl J, Singer M, Kubisch R, Schreiner L, Abhari BA, Wagner E, Kazmaier U, Fulda S, et al (2014) Pretubulysin: a new option for the treatment of metastatic cancer. Cell Death Dis 5: e1001–e1001

Burakov AV & Nadezhdina ES (2013) Association of nucleus and centrosome: magnet or velcro? Cell Biol Int 37: 95–104

Carden S, Vitiello E, Silva IR e, Holder J, Quarantotti V, Kishore K, Franklin VNR, D’Santos C, Ochi T, Breugel M van, et al (2023) Proteomic profiling of centrosomes across multiple mammalian cell and tissue types by an affinity capture method. Dev Cell 58: 2393–2410.e9

Castiel A, Visochek L, Mittelman L, Dantzer F, Izraeli S & Cohen-Armon M (2011) A phenanthrene derived PARP inhibitor is an extra-centrosomes de-clustering agent exclusively eradicating human cancer cells. BMC Cancer 11: 412

Chatterjee S, Sarkar A, Zhu J, Khodjakov A, Mogilner A & Paul R (2020) Mechanics of Multicentrosomal Clustering in Bipolar Mitotic Spindles. Biophys J 119: 434–447

Chauveau A, Aucher A, Eissmann P, Vivier E & Davis DM (2010) Membrane nanotubes facilitate long-distance interactions between natural killer cells and target cells. Proc Natl Acad Sci 107: 5545– 5550

Chiarugi A, Meli E, Calvani M, Picca R, Baronti R, Camaioni E, Costantino G, Marinozzi M, Pellegrini-Giampietro DE, Pellicciari R, et al (2003) Novel Isoquinolinone-Derived Inhibitors of Poly(ADP-ribose) Polymerase-1: Pharmacological Characterization and Neuroprotective Effects in an in Vitro Model of Cerebral Ischemia. J Pharmacol Exp Ther 305: 943–949

Combs J, Kim SJ, Tan S, Ligon LA, Holzbaur ELF, Kuhn J & Poenie M (2006) Recruitment of dynein to the Jurkat immunological synapse. Proc Natl Acad Sci 103: 14883–14888

Decarreau J, Wagenbach M, Lynch E, Halpern AR, Vaughan JC, Kollman J & Wordeman L (2017) The tetrameric kinesin Kif25 suppresses pre-mitotic centrosome separation to establish proper spindle orientation. Nat Cell Biol 19: 384–390

Dixit R & Cyr R (2004) Encounters between Dynamic Cortical Microtubules Promote Ordering of the Cortical Array through Angle-Dependent Modifications of Microtubule Behavior. Plant Cell 16: 3274–3284

Faragher AJ & Fry AM (2003) Nek2A kinase stimulates centrosome disjunction and is required for formation of bipolar mitotic spindles. Mol Biol Cell 14: 2876–2889

Firdous F, Raza HG, Chotana GA, Choudhary MI, Faisal A & Saleem RSZ (2023) Centrosome Clustering & Chemotherapy. Mini-Rev Med Chem 23: 429–451

Fırat-Karalar EN & Stearns T (2014) The centriole duplication cycle. Philos Trans R Soc B: Biol Sci 369: 20130460

Foethke D, Makushok T, Brunner D & Nédélec F (2009) Force- and length-dependent catastrophe activities explain interphase microtubule organization in fission yeast. Mol Syst Biol 5: 241–241

Fourriere L, Jimenez AJ, Perez F & Boncompain G (2020) The role of microtubules in secretory protein transport. J Cell Sci 133: jcs237016

Fourriere L, Kasri A, Gareil N, Bardin S, Bousquet H, Pereira D, Perez F, Goud B, Boncompain G & Miserey-Lenkei S (2019) RAB6 and microtubules restrict protein secretion to focal adhesions. J Cell Biol 218: 2215–2231

Fry AM, Mayor T, Meraldi P, Stierhof Y-D, Tanaka K & Nigg EA (1998) C-Nap1, a Novel Centrosomal Coiled-Coil Protein and Candidate Substrate of the Cell Cycle–regulated Protein Kinase Nek2. J Cell Biol 141: 1563–1574

Ganem NJ, Godinho SA & Pellman D (2009) A mechanism linking extra centrosomes to chromosomal instability. Nature 460: 278–282

Gorilak P, Pružincová M, Vachova H, Olšinová M, Cernohorska MS & Varga V (2021) Expansion microscopy facilitates quantitative super-resolution studies of cytoskeletal structures in kinetoplastid parasites. Open Biol 11: 210131

Grakoui A, Bromley SK, Sumen C, Davis MM, Shaw AS, Allen PM & Dustin ML (1999) The Immunological Synapse: A Molecular Machine Controlling T Cell Activation. Science 285: 221– 227

Grill SW & Hyman AA (2005) Spindle Positioning by Cortical Pulling Forces. Dev Cell 8: 461–465

Gunzer M, Weishaupt C, Hillmer A, Basoglu Y, Friedl P, Dittmar KE, Kolanus W, Varga G & Grabbe S (2004) A spectrum of biophysical interaction modes between T cells and different antigen-presenting cells during priming in 3-D collagen and in vivo. Blood 104: 2801–2809

Heald R & Khodjakov A (2015) Thirty years of search and capture: The complex simplicity of mitotic spindle assembly. J Cell Biol 211: 1103–1111

Hill TL (1985) Theoretical problems related to the attachment of microtubules to kinetochores. Proc Natl Acad Sci 82: 4404–4408

Holy TE & Leibler S (1994) Dynamic instability of microtubules as an efficient way to search in space. Proc Natl Acad Sci 91: 5682–5685

Horňák P, Kottfer D, Kyzioł K, Trebuňová M, Majerníková J, Kaczmarek Ł, Trebuňa J, Hašuľ J & Paľo M (2020) Microstructure and Mechanical Properties of Annealed WC/C PECVD Coatings Deposited Using Hexacarbonyl of W with Different Gases. Materials 13: 3576

Kopf A & Kiermaier E (2021) Dynamic Microtubule Arrays in Leukocytes and Their Role in Cell Migration and Immune Synapse Formation. Front Cell Dev Biol 9: 635511

Krämer A, Neben K & Ho AD (2005) Centrosome aberrations in hematological malignancies. Cell Biol Int 29: 375–383

Kuhn JR & Poenie M (2002) Dynamic Polarization of the Microtubule Cytoskeleton during CTL-Mediated Killing. Immunity 16: 111–121

Kupfer A & Dennert G (1984) Reorientation of the microtubule-organizing center and the Golgi apparatus in cloned cytotoxic lymphocytes triggered by binding to lysable target cells. J Immunol (Baltim, Md : 1950) 133: 2762–6

Kupfer A, Dennert G & Singer SJ (1985) The reorientation of the Golgi apparatus and the microtubule-organizing center in the cytotoxic effector cell is a prerequisite in the lysis of bound target cells. J Mol Cell Immunol : JMCI 2: 37–49

Kupfer A & Singer SJ (1989) The specific interaction of helper T cells and antigen-presenting B cells. IV. Membrane and cytoskeletal reorganizations in the bound T cell as a function of antigen dose. J Exp Med 170: 1697–1713

Kupfer A, Singer SJ & Dennert G (1986) On the mechanism of unidirectional killing in mixtures of two cytotoxic T lymphocytes. Unidirectional polarization of cytoplasmic organelles and the membrane-associated cytoskeleton in the effector cell. J Exp Med 163: 489–498

Kupfer A, Swain SL & Singer SJ (1987) The specific direct interaction of helper T cells and antigen-presenting B cells. II. Reorientation of the microtubule organizing center and reorganization of the membrane-associated cytoskeleton inside the bound helper T cells. J Exp Med 165: 1565–1580

Laan L, Pavin N, Husson J, Romet-Lemonne G, van Duijn M, López MP, Vale RD, Jülicher F, Reck-Peterson SL & Dogterom M (2012) Cortical Dynein Controls Microtubule Dynamics to Generate Pulling Forces that Position Microtubule Asters. Cell 148: 502–514

Laporte MH, Klena N, Hamel V & Guichard P (2022) Visualizing the native cellular organization by coupling cryofixation with expansion microscopy (Cryo-ExM). Nat Methods 19: 216–222

Leithner A, Altenburger LM, Hauschild R, Assen FP, Rottner K, Stradal TEB, Diz-Muñoz A, Stein JV & Sixt M (2021) Dendritic cell actin dynamics control contact duration and priming efficiency at the immunological synapse. J Cell Biol 220: e202006081

Letort G, Nedelec F, Blanchoin L & Théry M (2016) Centrosome centering and decentering by microtubule network rearrangement. Mol Biol Cell 27: 2833–2843

Li R & Gundersen GG (2008) Beyond polymer polarity: how the cytoskeleton builds a polarized cell. Nat Rev Mol Cell Biol 9: 860–873

Liu Z-G, Smith SW, McLaughlin KA, Schwartz LM & Osborne BA (1994) Apoptotic signals delivered through the T-cell receptor of a T-cell hybrid require the immediate–early gene nur77. Nature 367: 281–284

Luxton GG & Gundersen GG (2011) Orientation and function of the nuclear–centrosomal axis during cell migration. Curr Opin Cell Biol 23: 579–588

Mallick A, Sarkar A & Paul R (2022) A force-balance model for centrosome positioning and spindle elongation during interphase and anaphase B. Indian J Phys 96: 2667–2691

Mardin BR, Lange C, Baxter JE, Hardy T, Scholz SR, Fry AM & Schiebel E (2010) Components of the Hippo pathway cooperate with Nek2 kinase to regulate centrosome disjunction. Nat Cell Biol 12: 1166–1176

Martín-Cófreces NB, Baixauli F & Sánchez-Madrid F (2014) Immune synapse: conductor of orchestrated organelle movement. Trends Cell Biol 24: 61–72

Martín-Cófreces NB, Robles-Valero J, Cabrero JR, Mittelbrunn M, Gordón-Alonso M, Sung C-H, Alarcón B, Vázquez J & Sánchez-Madrid F (2008) MTOC translocation modulates IS formation and controls sustained T cell signaling. J Cell Biol 182: 951–962

Meyer-Gerards C & Bazzi H (2024) Developmental and tissue-specific roles of mammalian centrosomes. FEBS J

Mittelbrunn M, Hoyo GM del, López-Bravo M, Martín-Cofreces NB, Scholer A, Hugues S, Fetler L, Amigorena S, Ardavín C & Sánchez-Madrid F (2009) Imaging of plasmacytoid dendritic cell interactions with T cells. Blood 113: 75–84

Möller K, Brambach M, Villani A, Gallo E, Gilmour D & Peri F (2022) A role for the centrosome in regulating the rate of neuronal efferocytosis by microglia in vivo. eLife 11: e82094

Monks CRF, Freiberg BA, Kupfer H, Sciaky N & Kupfer A (1998) Three-dimensional segregation of supramolecular activation clusters in T cells. Nature 395: 82–86

Moran AE, Holzapfel KL, Xing Y, Cunningham NR, Maltzman JS, Punt J & Hogquist KA (2011) T cell receptor signal strength in Treg and iNKT cell development demonstrated by a novel fluorescent reporter mouse. J Exp Med 208: 1279–1289

Muroyama A & Lechler T (2017) Microtubule organization, dynamics and functions in differentiated cells. Development 144: 3012–3021

Nigg EA & Holland AJ (2018) Once and only once: mechanisms of centriole duplication and their deregulation in disease. Nat Rev Mol Cell Biol 19: 297–312

Oakley C & Brunette DM (1993) The sequence of alignment of microtubules, focal contacts and actin filaments in fibroblasts spreading on smooth and grooved titanium substrata. J Cell Sci 106: 343– 354

Paintrand M, Moudjou M, Delacroix H & Bornens M (1992) Centrosome organization and centriole architecture: Their sensitivity to divalent cations. J Struct Biol 108: 107–128

Paul R, Wollman R, Silkworth WT, Nardi IK, Cimini D & Mogilner A (2009) Computer simulations predict that chromosome movements and rotations accelerate mitotic spindle assembly without compromising accuracy. Proc Natl Acad Sci 106: 15708–15713

Pavin N & Tolić-Nørrelykke IM (2014) Swinging a sword: how microtubules search for their targets. Syst Synth Biol 8: 179–186

Pereira SG, Louro MAD & Bettencourt-Dias M (2021) Biophysical and Quantitative Principles of Centrosome Biogenesis and Structure. Annu Rev Cell Dev Biol 37: 1–21

Picone R, Ren X, Ivanovitch KD, Clarke JDW, McKendry RA & Baum B (2010) A Polarised Population of Dynamic Microtubules Mediates Homeostatic Length Control in Animal Cells. PLoS Biol 8: e1000542

Pihan GA, Purohit A, Wallace J, Knecht H, Woda B, Quesenberry P & Doxsey SJ (1998) Centrosome defects and genetic instability in malignant tumors. Cancer Res 58: 3974–85

Pihan GA, Wallace J, Zhou Y & Doxsey SJ (2003) Centrosome abnormalities and chromosome instability occur together in pre-invasive carcinomas. Cancer Res 63: 1398–404

Prosser SL & Pelletier L (2017) Mitotic spindle assembly in animal cells: a fine balancing act. Nat Rev Mol Cell Biol 18: 187–201

Pulecio J, Petrovic J, Prete F, Chiaruttini G, Lennon-Dumenil A-M, Desdouets C, Gasman S, Burrone OR & Benvenuti F (2010) Cdc42-mediated MTOC polarization in dendritic cells controls targeted delivery of cytokines at the immune synapse. J Exp Med 207: 2719–2732

Quah BJC, Warren HS & Parish CR (2007) Monitoring lymphocyte proliferation in vitro and in vivo with the intracellular fluorescent dye carboxyfluorescein diacetate succinimidyl ester. Nat Protoc 2: 2049–2056

Sarkar A, Rieger H & Paul R (2019) Search and Capture Efficiency of Dynamic Microtubules for Centrosome Relocation during IS Formation. Biophys J 116: 2079–2091

Sato N, Mizumoto K, Nakamura M, Nakamura K, Kusumoto M, Niiyama H, Ogawa T & Tanaka M (1999) Centrosome abnormalities in pancreatic ductal carcinoma. Clin cancer Res : Off J Am Assoc Cancer Res 5: 963–70

Shaw AS & Dustin ML (1997) Making the T Cell Receptor Go the Distance: A Topological View of T Cell Activation. Immunity 6: 361–369

Som S, Chatterjee S & Paul R (2019) Mechanistic three-dimensional model to study centrosome positioning in the interphase cell. Phys Rev E 99: 012409

Som S & Paul R (2023) Mechanistic model for nuclear migration in hyphae during mitosis. Phys Rev E 108: 014401

Steinman RM & Cohn ZA (1973) IDENTIFICATION OF A NOVEL CELL TYPE IN PERIPHERAL LYMPHOID ORGANS OF MICE. J Exp Med 137: 1142–1162

Stinchcombe JC, Majorovits E, Bossi G, Fuller S & Griffiths GM (2006) Centrosome polarization delivers secretory granules to the immunological synapse. Nature 443: 462–465

Tamzalit F, Tran D, Jin W, Boyko V, Bazzi H, Kepecs A, Kam LC, Anderson KV & Huse M (2020) Centrioles control the capacity, but not the specificity, of cytotoxic T cell killing. Proc Natl Acad Sci 117: 4310–4319

Ueda H, Morphew MK, McIntosh JR & Davis MM (2011) CD4+ T-cell synapses involve multiple distinct stages. Proc Natl Acad Sci 108: 17099–17104

Ullrich A, Chai Y, Pistorius D, Elnakady YA, Herrmann JE, Weissman KJ, Kazmaier U & Müller R (2009) Pretubulysin, a Potent and Chemically Accessible Tubulysin Precursor from Angiococcus disciformis. Angew Chem Int Ed 48: 4422–4425

Varga V, Leduc C, Bormuth V, Diez S & Howard J (2009) Kinesin-8 Motors Act Cooperatively to Mediate Length-Dependent Microtubule Depolymerization. Cell 138: 1174–1183

Waterman-Storer CM & Salmon ED (1997) Actomyosin-based Retrograde Flow of Microtubules in the Lamella of Migrating Epithelial Cells Influences Microtubule Dynamic Instability and Turnover and Is Associated with Microtubule Breakage and Treadmilling. J Cell Biol 139: 417–434

Weier A-K, Homrich M, Ebbinghaus S, Juda P, Miková E, Hauschild R, Zhang L, Quast T, Mass E, Schlitzer A, et al (2022) Multiple centrosomes enhance migration and immune cell effector functions of mature dendritic cells. J Cell Biol 221: e202107134

Wollman R, Cytrynbaum EN, Jones JT, Meyer T, Scholey JM & Mogilner A (2005) Efficient Chromosome Capture Requires a Bias in the ‘Search-and-Capture’ Process during Mitotic-Spindle Assembly. Curr Biol 15: 828–832

Wong YL, Anzola JV, Davis RL, Yoon M, Motamedi A, Kroll A, Seo CP, Hsia JE, Kim SK, Mitchell JW, et al (2015) Reversible centriole depletion with an inhibitor of Polo-like kinase 4. Science 348: 1155–1160

Wu X, Xiang X & Hammer JA (2006) Motor proteins at the microtubule plus-end. Trends Cell Biol 16: 135–143

Yi J, Wu X, Chung AH, Chen JK, Kapoor TM & Hammer JA (2013) Centrosome repositioning in T cells is biphasic and driven by microtubule end-on capture-shrinkage. J Cell Biol 202: 779–792

Yuseff M-I, Reversat A, Lankar D, Diaz J, Fanget I, Pierobon P, Randrian V, Larochette N, Vascotto F, Desdouets C, et al (2011) Polarized Secretion of Lysosomes at the B Cell Synapse Couples Antigen Extraction to Processing and Presentation. Immunity 35: 361–374

Zhu J, Burakov A, Rodionov V & Mogilner A (2010) Finding the Cell Center by a Balance of Dynein and Myosin Pulling and Microtubule Pushing: A Computational Study. Mol Biol Cell 21: 4418–4427

