## Appendix for "Multiple clustered centrosomes in antigen-presenting cells foster T cell activation without MTOC polarization"

**Extended view figures**

**Supplementary methods**

**Table S1**

**Appendix references**

Extended View Figures

### Extended View Figure EV1

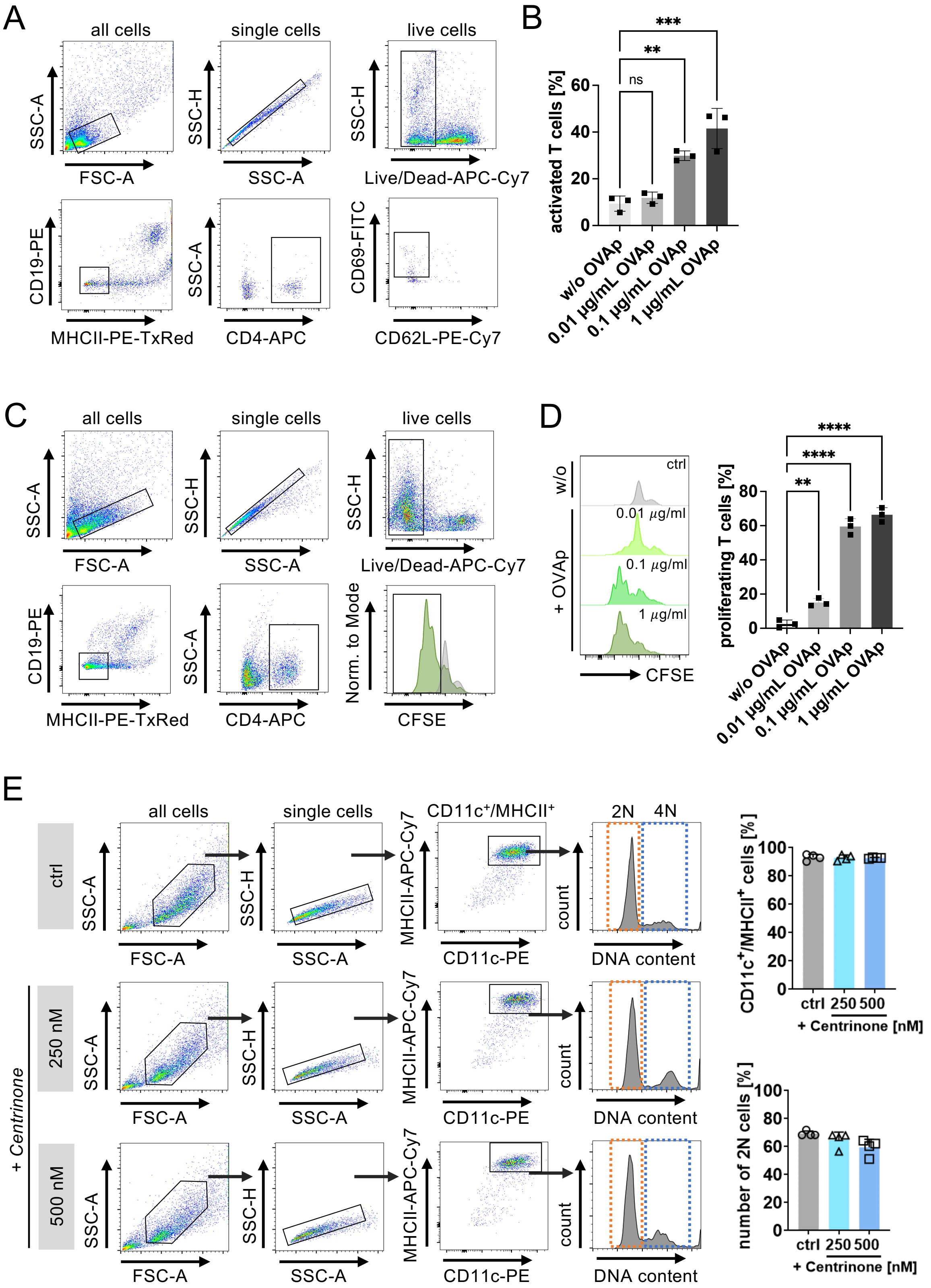

**Fig. EV1.**

**An intact centrosome in APCs is required for efficient T cell activation.** (**A**) Gating strategy for analysing $CD69^+/CD62L^-$ activated $CD4^+$ T cells in the absence of antigen (w/o OVAp) or in the presence of different concentrations of OVAp. (**B**) Quantification of antigen-specific T cell activation. Graph shows mean values ± s.d. of three technical replicates of one out of four independent experiments. **, $P < 0.0021$; ***, $P < 0.001$ (one-way Anova with Dunnett's multiple comparisons). (**C**) Gating strategy for quantifying T cell proliferation in the absence of antigen (w/o OVAp) or in the presence of different concentrations of OVAp. Unstained samples served as control and were included as light grey filled line. (**D**) Quantification of antigen-specific T cell proliferation. Graph shows mean values ± s.d. of three technical replicates of one out of four independent experiments. **, $P < 0.0021$; ****, $P < 0.0001$ (one-way Anova with Dunnett's multiple comparisons). (**E**) Differentiation and maturation of BMDCs in the presence of the PLK4 inhibitor Centrinone. Left: gating strategy to assess DC differentiation in Centrinone-treated and control cells. Mature DCs were identified as $MHCII^+/CD11c^+$ cells and further analyzed for DNA content. Right: quantification of $MHCII^+/CD11c^+$ (upper graph) and 2N (lower graph) cells in Centrinone-treated and control cells. Graphs show mean values ± s.d. of 4 independent experiments with cells derived from 4 different mice. (A-D) $N = 10.000$ cells per condition.

### Extended View Figure EV2

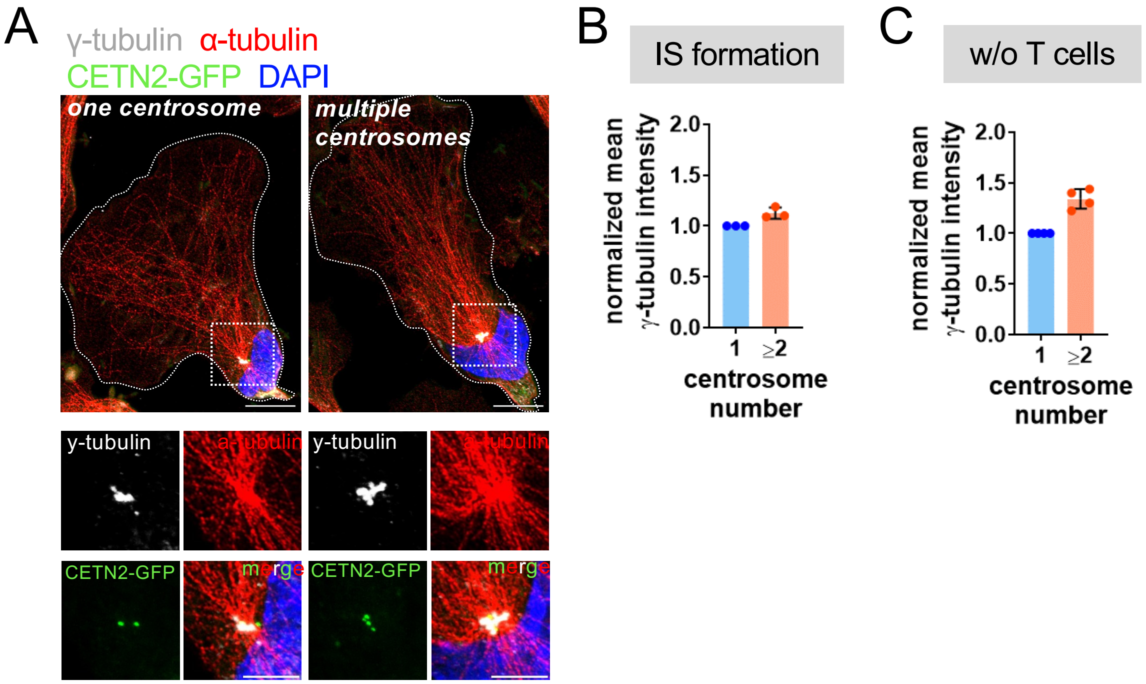


**Fig. EV2.**

**IS formation in the presence of multiple centrosomes.** (**A**) Immunostaining of MT filaments in sorted 2N mature CETN2-GFP expressing BMDCs without T cells. White dotted boxes indicate magnified regions at the bottom. Merged and individual channels of CETN2-GFP (green), γ-tubulin (white) and α-tubulin (red) are shown. Cells were counterstained with DAPI (blue). Scale bars, 10 $\mu$m (upper panels) and 2 $\mu$m (insets bottom). (**B**) (**C**) Quantification of γ-tubulin signal intensity in DCs in the presence (**B**) or absence (**C**) of T cells. Values for one centrosome were set to one. Each data point represents one independent experiment with at least $N = 100$ cells analyzed per experiment.

### Extended View Figure EV3

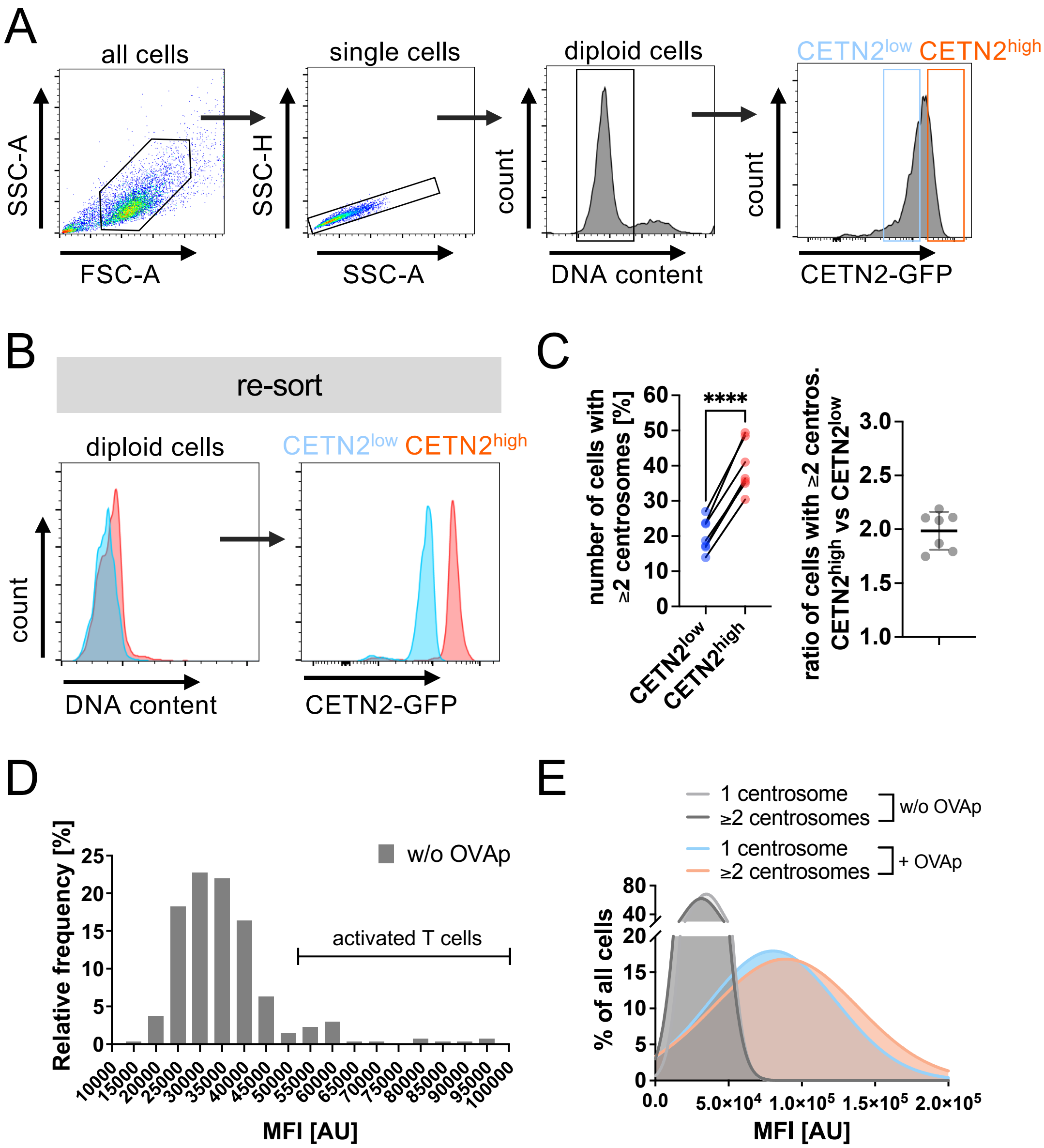


**Fig. EV3.**

**Enhanced T cell activation in the presence of multiple centrosomes. (A)** Separation of CETN2-GFP$^{low}$ (blue) and CETN2-GFP$^{high}$ (red) expressing mature BMDCs. Single cells were gated on 2N and further separated according to CETN2-GFP signal intensities into CETN2-GFP high and low expressing cells. (**B**) Post-sort analysis of CETN2-GFP$^{low}$ (blue) and CETN2-GFP$^{high}$ (red) expressing cells for DNA content (left) and CETN2-GFP signal intensities (right). (**C**) Left: quantification of percentage of cells with ≥2 centrosomes in CETN2-GFP$^{low}$ (blue) and CETN2-GFP$^{high}$ (red) expressing cells. Centrosome numbers of sorted DC subpopulations were determined by confocal microscopy according to CETN2-GFP/$\gamma$-tubulin$^{+}$

foci. Each data point represents one independent experiment. ****, $P < 0.0001$ (two-tailed, paired Student's *t*-test). Right graph: ratio of cells with multiple centrosomes between CETN2-GFP$^{high}$ and CETN2-GFP$^{low}$ cells. Frequency distribution of mean fluorescence intensities of Nur77$^{GFP}$ expression levels in the absence (**D**) or presence (**E**) of OVAp and in dependence of centrosome numbers (**E**). Threshold for T cell activation was set according to w/o OVAp sample and is indicated in D.

### Extended View Figure EV4

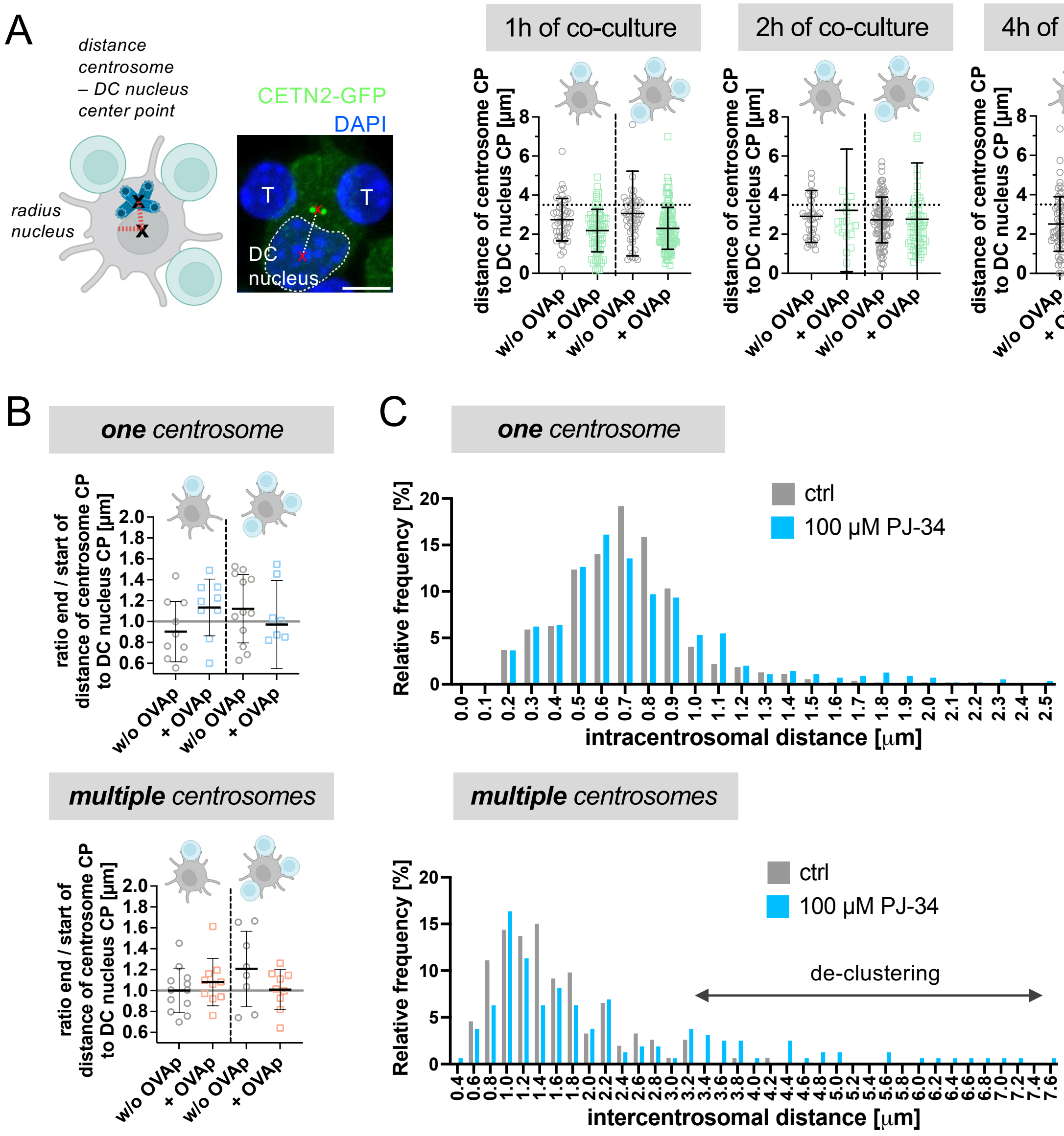


**Fig. EV4.**

**Centrosome configuration during antigen-specific DC-T cell contacts.** (**A**) Quantification of centrosome positioning relative to the nucleus. Left: sketch indicating distance of centrosome to center of nucleus and radius of the nucleus (red line). Scale bar, 5 $\mu$m. Right: quantification of distance of centrosome to CP of the nucleus. Graphs represent mean values ± s.d.. Each data point depicts one cell derived from 3 independent experiments. Dotted lines indicate radius of the nucleus. (**B**) Quantification of ratio of distances between centrosome(s) CP and the center of the DC nucleus at the end vs. the beginning of recording. Graphs represent mean values ± s.d. Each data point depicts one cell derived from 5 independent experiments. (**C**) Frequency distribution of intracentrosomal (upper) and intercentrosomal (lower) distances in mature CETN2-GFP expressing DCs after PJ-34 treatment and control cells. Shift of intracentrosomal distances to larger values indicates perturbed centrosome linker integrity; larger intercentrosomal distances denote de-clustering of multiple centrosomes.

#### Extended View Figure EV5

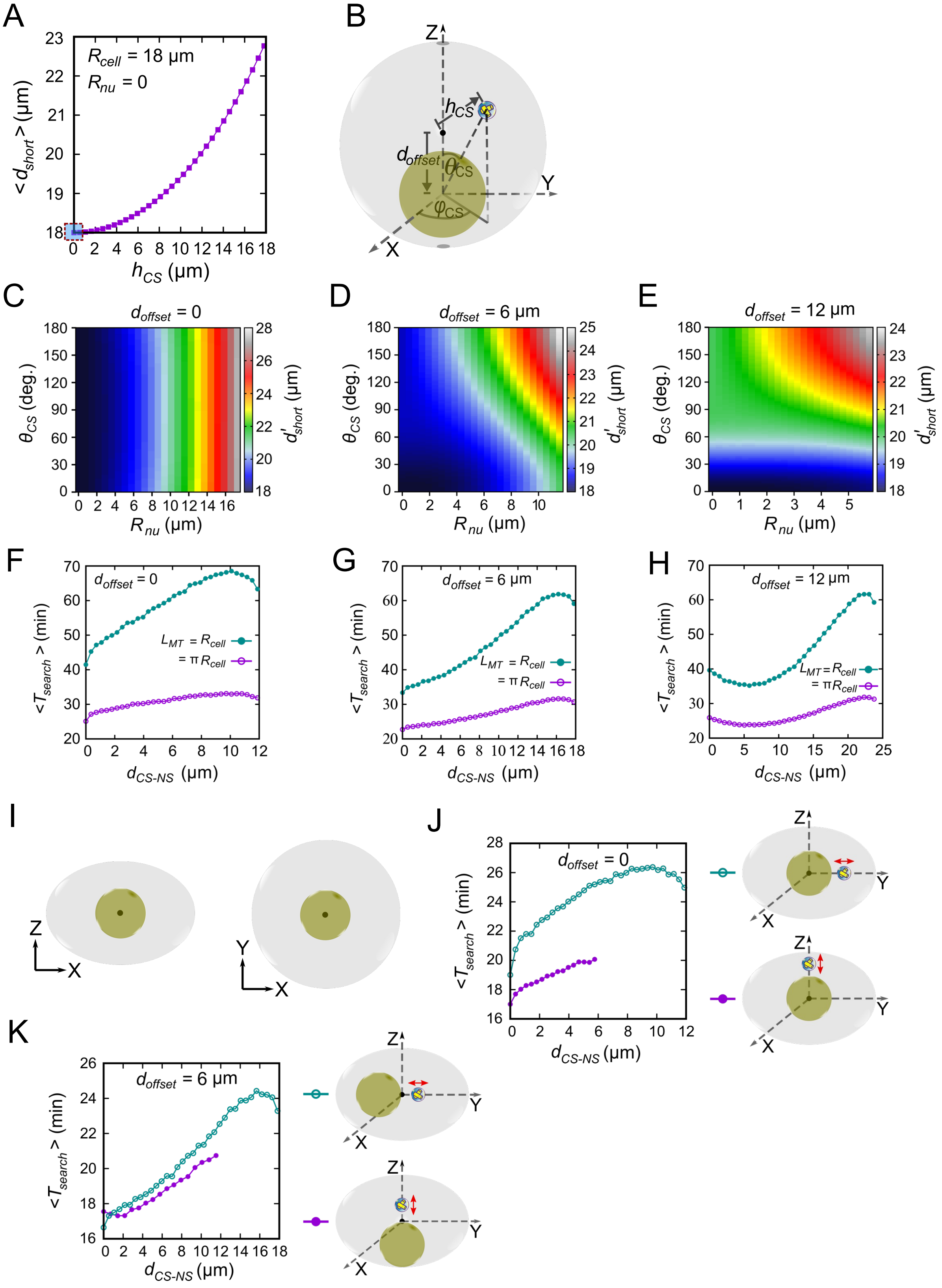

**Fig. EV5.**

**Modelling optimal centrosome position and configuration in DCs.** (**A**) Geometrically optimal centrosome position in the absence of a nucleus. Plot of the average shortest distance between the centrosome and the target points on the cell surface, $< d_{short} >$, as a function of the centrosome distance from the cell center, $h_{CS}$. The red outlined box denotes the optimized value of $< d_{short} >$ when the centrosome is placed at the cell center, as obtained from both simulations and our analytical calculations in *Supplementary Materials.* (**B-E**) Centrosome placed along the line joining the cell and nucleus center gives the global minimum in the average shortest distance between the centrosome and the target points on the cell surface. (**B**) A schematic depiction of the centrosome position denoted by the polar angle $\theta_{CS}$ and the azimuthal angle $\phi_{CS}$. The values of $< d_{short} >$ involving isotropic target points on the cell surface do not depend on $\phi_{CS}$, due to the spherical symmetry of both the cell and the nucleus in the model. (**C-E**) Values of $d'_{short}$ plotted as a function of $\theta_{CS}$ and nuclear radius, $R_{nu}$, for different off-centered positions of the nucleus. $d'_{short}$ represents the optimal value of $< d_{short} >$ obtained by varying the centrosome's position at different distances from the nucleus while keeping the $\theta_{CS}$ fixed. **(F-H)** Average search times dependent on average MT length. The average search time, $< T_{search} >$, varies with centrosome position away from nuclear surface, $d_{CS-NS}$ , showing qualitatively similar behavior for different average MT lengths, albeit with comparatively smaller search time for MTs having larger average length. (**I-K**) Optimal centrosome positioning in flattened cells. The cell is flattened along the z-axis of the cell, which is also reflected from the schematic depiction of the corresponding XZ and XY views of the cell (**I**). The semi-axes of the cell are $A_x = 18$ µm, $A_y = 18$ µm, and $A_z = 12$ µm, respectively. (**J**) $< T_{search} >$ vs $d_{CS-NS}$ plot for a centrally located nucleus ($d_{offet} = 0$) demonstrating a similar optimal perinuclear centrosome positioning as in rounded spherical cells, but with the centrosome situated along the short axis of the flattened DCs. (**K**) Similar plot but for an off-centered nucleus ($d_{offet} = 6$ µm). Like rounded cells, the plot indicates optimal centrosome positioning at the cell center. The search time is smaller when the nucleus is shifted along the long axis of the cell compared to when it is shifted along the short axis. (**J-K**) The double-headed red arrow denotes the direction of centrosome shifting along the short and long axes of the cell, respectively. The MTs are not allowed to glide along the cell surface.

### Extended View Figure EV6

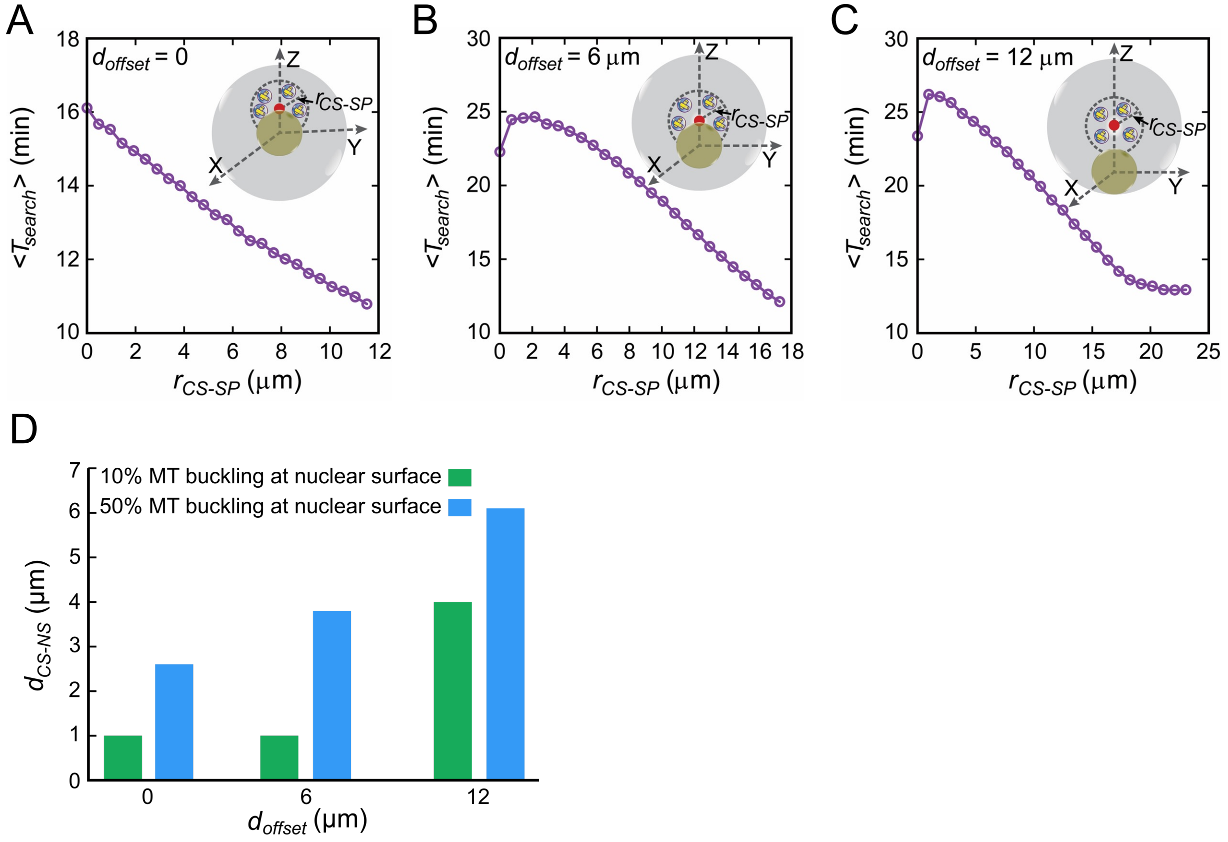


**Fig. EV6.**

**Model predictions of search time with dispersed centrosomes and freely gliding MTs along the cell surface, and a mechanistic model regulating the positioning of closely placed centrosomes.**

**(A-C)** Dispersed centrosomes reduce the search time in the presence of unrestricted MTs gliding along the cell surface. Average search time, $< T_{search} >$, is plotted against $r_{CS-SP}$ for different off-centered positions of the nucleus. $r_{CS-SP}$ is the radius of the imaginary sphere centered around a specifically chosen point (red) inside the cell (the points are considered perinuclear for $d_{offset}$= 0 and at the cell center for $d_{offset}$= 6 μm, and 12 μm), within which centrosomes are randomly placed, as shown schematically in the insets of the figures. (**D**) The mechanistic force-balance model demonstrates the rise in the distance between the centrosome cluster and the nuclear surface when a higher fraction of MTs is assumed to buckle at the nuclear surface. The data in the plot is compared for scenarios where either 10% or 50% of the MTs hitting the nucleus buckle at the nuclear surface.

**Movie legend**

**Movie EV1**

**Live cell imaging of centrosome dynamics during immune synapse formation**

Time-lapse live-cell confocal microscopy of antigen-specific DC-T cell contacts. Merged channels of CETN2-GFP (green), $Ca^{2+}$-Cal520 (green) and DNA stain (Vybrant Dye Cycle Violet, blue) are displayed. Movie shows DC carrying one centrosome and forming a mono-conjugated synapse. White arrow heads point to centrosome(s) position. Cell-cell contacts were imaged with frame rate of 20 s. Played as 10 frames/s. Scalebar 10 µm.

**Movie EV2**

**Live cell imaging of centrosome dynamics during immune synapse formation**

Time-lapse live-cell confocal microscopy of antigen-specific DC-T cell contacts. Merged channels of CETN2-GFP (green), $Ca^{2+}$-Cal520 (green) and DNA stain (Vybrant Dye Cycle Violet, blue) are displayed. Movie shows DC carrying one centrosome and forming a multi-conjugated synapse. White arrow heads point to centrosome(s) position. Cell-cell contacts were imaged with frame rate of 20 s. Played as 10 frames/s. Scalebar 10 µm.

**Movie EV3**

**Live cell imaging of centrosome dynamics during immune synapse formation**

Time-lapse live-cell confocal microscopy of antigen-specific DC-T cell contacts. Merged channels of CETN2-GFP (green), $Ca^{2+}$-Cal520 (green) and DNA stain (Vybrant Dye Cycle Violet, blue) are displayed. Movie shows DC carrying multiple centrosomes and forming a mono-conjugated synapse. White arrow heads point to centrosome(s) position. Cell-cell contacts were imaged with frame rate of 20 s. Played as 10 frames/s. Scalebar 10 µm.

**Movie EV4**

**Live cell imaging of centrosome dynamics during immune synapse formation**

Time-lapse live-cell confocal microscopy of antigen-specific DC-T cell contacts. Merged channels of CETN2-GFP (green), $Ca^{2+}$-Cal520 (green) and DNA stain (Vybrant Dye Cycle Violet, blue) are displayed. Movie shows DC carrying multiple centrosomes and forming a multi-conjugated synapse. White arrow heads point to centrosome(s) position. Cell-cell contacts were imaged with frame rate of 20 s. Played as 10 frames/s. Scalebar 10 µm.

**Movie EV5**

Simulation movie demonstrating the dynamics of four closely placed centrosomes with a nucleus fixed at the cell center ($d_{offset} = 0$). The cell is depicted in gray, the MTs in blue, the nucleus in olive, and the centrosomes are shown in yellow.

**Movie EV6**

Simulation movie demonstrating the dynamics of four closely placed centrosomes when the nucleus is off-centered such that the nuclear surface touches the cell center ($d_{offset} = 6$ µm). The cell is depicted in gray, the MTs in blue, the nucleus in olive, and the centrosomes are shown in yellow.Movie EV7

Simulation movie demonstrating the dynamics of four closely placed centrosomes when the nucleus is highly off-centered such that the nuclear surface touches the cell surface ($d_{offset} = 12$ µm). The cell is depicted in gray, the MTs in blue, the nucleus in olive, and the centrosomes are shown in yellow.

**Movie EV7**

Simulation movie demonstrating the dynamics of four closely placed centrosomes when the nucleus is highly off-centered such that the nuclear surface touches the cell surface ($d_{offset} = 12$ µm). The cell is depicted in gray, the MTs in blue, the nucleus in olive, and the centrosomes are shown in yellow.

**Supplementary methods**

**Analytical prediction of optimal centrosome position in the absence of a nucleus**

We find the optimal centrosome position, corresponding to the minimum average geometric distance between the centrosome and the cell surface points, inside a 2*D* circular and 3*D* spherical cell of radius $R_{cell}$ without a nucleus.

**In** 2*D* **circular cell:** In the Cartesian coordinate frame, consider the position of centrosome $(x_0, y_0)$ and the cell center at the origin $(0, 0)$. Then, the distance between an arbitrary point $(x, y)$ on the cell surface and the centrosome is

$$d^2 = (x - x_0)^2 + (y - y_0)^2$$

$$= (R_{cell}cos\theta - x_0)^2 + (R_{cell}sin\theta - y_0)^2$$

Here, $R_{cell}cos\theta$ and $R_{cell}sin\theta$ represent the parametric equations of $x$ and $y$, respectively, in the plane polar coordinate system. $\theta$ $(\in 0 - 360^0)$ is the counterclockwise angle (also called polar angle) measured with respect to the positive x-axis of the cell. Therefore, the average distance relative to the entire cell surface would be

$$< d^2 > = \frac{1}{2\pi}\int_0^{2\pi}(R_{cell}cos\theta - x_0)^2 + \ (R_{cell}sin\theta - y_0)^2 \, d\varphi$$

$$= \frac{1}{2\pi}\int_0^{2\pi} d\varphi \left\{R_{cell}^2(cos^2\theta \ + \ sin^2\theta) - 2R_{cell}x_0cos\varphi - 2R_{cell}y_0sin\varphi + (x_0^2 \ + \ y_0^2)\right\}$$

$$= R_{cell}^2 + x_0^2 \ + \ y_0^2$$

Therefore,

$$< d^2 >_{min} = R_{cell}^2 \text{ with } x_{0,min} = 0, y_{0,min} = 0$$

$$< d^2 >_{max} = 2R_{cell}^2 \text{ with } x_{0,max}, \ y_{0,max} \in \text{points on the cell surface}$$

Therefore, the minimum of the average geometric distance, $< d_{short} >$ $(= \sqrt{< d^2 >_{min}} = R_{cell}$ ), correspond to the optimal centrosome position at the center of the circular cell.

**In** 3*D* **spherical cell:** Similar to above, the distance between an arbitrary 3*D* point $(x, y, z)$ on the surface of a spherical cell (center is at $(0, 0, 0)$) and the centrosome location $(x_0, y_0, z_0)$ follows

$$d^2 = (x - x_0)^2 + (y - y_0)^2 + (z - z_0)^2$$

$$= (R_{cell}sin\theta cos\varphi - x_0)^2 + (R_{cell}sin\theta sin\varphi - y_0)^2 + (R_{cell}cos\theta - z_0)^2$$

Here, $R_{cell}sin\theta cos\varphi$, $R_{cell}sin\theta sin\varphi$, and $R_{cell}cos\theta$ represent the parametric equation of $x$, $y$, and $z$, respectively, in a spherical polar coordinate system. $\theta$ ($\in 0-180^0$) and $\varphi$ ($\in 0-360^0$) denote the polar and azimuthal angle, measured with respect to the positive $z$ and $x$ axes of the cell, respectively.
Therefore, the average distance relative to the entire cell surface would be

$$< d^2 > = \frac{1}{4\pi}\int_0^{2\pi} d\varphi \int_0^{\pi} d\theta sin\theta \ (x-x_0)^2 + (y-y_0)^2 + (z-z_0)^2$$

$$= \frac{1}{4\pi}\int_0^{2\pi} d\varphi \int_0^{\pi} d\theta sin\theta \ \{(x^2+y^2+z^2) + (x_0^2+y_0^2+z_0^2) - 2(xx_0+yy_0+zz_0)\}$$

$$= R_{cell}^2 + (x_0^2+y_0^2+z_0^2) - \frac{1}{2\pi}\int_0^{2\pi} d\varphi \int_0^{\pi} d\theta sin\theta \ R_{cell}(x_0 sin\theta cos\varphi + y_0 sin\theta sin\varphi + z_0 cos\theta)$$

$$= R_{cell}^2 + x_0^2 + y_0^2 + z_0^2$$

Therefore,

$< d^2 >_{min} = R_{cell}^2$ with $x_{0,min} = 0, y_{0,min} = 0, z_{0,min} = 0$

$< d^2 >_{max} = 2R_{cell}^2$ with $x_{0,max}, y_{0,max}, z_{0,max} \in$ points on the cell surface

Therefore, the minimum of the average geometric distance, $< d_{short} >$ ($= \sqrt{< d^2 >_{min}} = R_{cell}$ ), correspond to the optimal centrosome position at the center of the spherical cell.

**Mathematical estimation of average search time in the absence of a nucleus predicts enhanced T cell priming capacity with increased microtubule numbers**

Efficient MT docking at the IS in T cells has recently been described by a mathematical model (*1*). The average search time of dynamic MTs in DCs can be estimated within this model framework. For a spherical cell of radius $R_{cell} = 18$ µm and without a nucleus, the average distance between the optimally localized centrosome at the cell center and an arbitrary point on the cell boundary is 18 µm. For a MT growing from the centrosome with a velocity $\sim 15$ µm/min (*2*) needs around 72 sec to reach the cell boundary without undergoing any catastrophe. In case the MT hits the IS it docks, in case it does not hit the IS it shrinks again with approximately the same velocity as it grows, which means that an unsuccessful growth attempt (i.e. not hitting the IS) needs approximately $\tau_{trial} = 144$ sec.

The probability, $p_{dock}$, for a single growing MT to dock at a single target IS, is equal to the ratio of the target (IS) area and the total cell area: $p_{dock} = \frac{A_\tau}{A_{cell}} = \frac{\pi R_\tau^2}{4\pi R_{cell}^2} \approx 0.0031$ for an IS radius of $R_\tau = 2$ µm (*3*). Consequently, a single MT needs on average $1/p_{dock} \approx 323$ search trials to dock at a single IS. In case of $n$ synapses, the number of trials is reduced by a factor of $1/n$ since the total area of $n$ synapses is simply $n$ times the area of one IS. DCs contain between $N_{MT} \approx 35$ and 45 dynamic MTs during IS formation performing simultaneous search. The probability that at least one of $N_{MT}$ simultaneously growing MTs docks at a single IS is derived as follows: the probability that a single growing MT does *not* dock at the IS is $(1 - p_{dock})$, hence the probability that $N_{MT}$ growing MTs do *not* dock at the IS is $(1 - p_{dock})^{N_{MT}}$, and therefore the probability that at least one MT docks is $1 - (1 - p_{dock})^{N_{MT}} = N_{MT}p_{dock} + \frac{1}{2} N_{MT}(N_{MT} - 1)p_{dock}^2 + \ldots$ . When $N_{MT}.p_{dock}$ is much smaller than 1, one can neglect the terms following $N_{MT}.p_{dock}$. Therefore, the probability that among $N_{MT}$ dynamic MTs performing a simultaneous search, at least one docks at the IS, is simply $N_{MT}$ times the probability that a single MT docks. Note that for large $N_{MT}$, i.e. when $N_{MT}p_{dock}$ is of order 1, one cannot neglect terms of higher order in $N_{MT}p_{dock}$ any more, the correct expression for the probability then being $1 - (1 - p_{dock})^{N_{MT}}$, which is still monotonously increasing with the number of MTs, $N_{MT}$.

The docking probability can immediately be translated into the average time that a dynamic MT needs to dock at a single IS: one unsuccessful growth-shrinkage trial lasts on average $\tau_{trial} = 144$ sec and on average a single MT needs $1/p_{dock} = 323$ trials to dock. Thus, a single MT needs on average $\tau_{trial} \cdot \frac{1}{p_{dock}} \approx 775$ min to dock at a single IS, and $N_{MT}$ MTs need $1/(N_{MT}\, p_{dock})$ trials, i.e. only a fraction of $1/N_{MT}$ of the time for a single MT: $35/40$ MTs need on average $\sim 22/19$ minutes to dock at a single IS, and $45/50$ MTs need $\sim 17/15$ min. An immediate prediction of these considerations is that the docking efficiency of MTs increases with increasing numbers of MTs performing search and capture dynamics, which in turn increases T cell priming capacity.

| | **Table S1: List of parameters** | |
|---|---|---|
| **Abbreviations** | **Meaning** | **Value \| Range \| Reference** |
| $R_{cell}$ | Cell radius | 18 μm \| (*4*) |
| $R_{nu}$ | Nucleus radius | $R_{cell}/3$ \| (*4*) |
| $d_{offset}$ | Distance between the nucleus center and cell center | 0, 6 μm, and 12 μm \| This study |
| $R_{\tau}$ | Target radius | 2 μm \| (*3*) |
| $N_{MT}$ | Total number of MTs | 40 \| (*4*) |
| $L_{MT}$ | Average MT length | $\pi R_{cell}$ \| $R_{cell} - \pi R_{cell}$ \| This study |
| $v_g$ | MTs growth velocity | 14.3 μm/min \| (*1*, *2*) |
| $v_s$ | MTs shrinkage velocity | 16 μm/min \| (*1*, *2*) |
| $f_c$ | MTs catastrophe frequency | $v_g/L_{MT}$ \| (*5*) |
| $f_r$ | MTs rescue frequency | 0 \| (*1*, *2*, *6*) |
| $k_0, \alpha_d$ | The phenomenological constant determining the sensitivity of dissociation of nuclear gliding MTs | 0.1, $\pi/2$ radian \| This study* |
| $\lambda_l$ | The phenomenological constant determining the increasing rate of catastrophe frequency of cell surface-gliding MTs | $0.1 - 10$ μm$^{-1}$ \| This study |
| $\eta$ | Co-efficient of cytoplasmic viscosity | ~200 pN s μm$^{-2}$ \| (*7*, *8*) |

**Table S1**

**Parameters used in the computational model to simulate the observed centrosomal arrangements in DCs.**

* The value of $\boldsymbol{k_0}$ and $\boldsymbol{\alpha_d}$ in Table S1 are chosen in a manner such that they ensure the successful capture of targets across a broad range of positions that are not directly visible to MTs from centrosome due to nuclear hindrance. A larger $\boldsymbol{k_0}$ values and/or smaller $\boldsymbol{\alpha_d}$ values may result in faster MT dissociation, potentially leaving targets positioned well below the equatorial plane (hiding far below the nucleus) uncaptured. Conversely, a smaller $\boldsymbol{k_0}$ values and/or larger $\boldsymbol{\alpha_d}$ values encourage MTs to continue gliding along the nuclear surface, delaying their dissociation. This can pose challenges when capturing targets that have just become directly inaccessible to MTs due to nuclear hindrance. Therefore, intermediate values of $\boldsymbol{k_0}$ and $\boldsymbol{\alpha_d}$ as used in Table S1 can ensure the target capture for all positioning of target for which the direct capture is not possible.

#### Appendix References

1. A. Sarkar, H. Rieger, R. Paul, Search and Capture Efficiency of Dynamic Microtubules for Centrosome Relocation during IS Formation. *Biophys. J.* **116**, 2079–2091 (2019).

2. T. E. Holy, S. Leibler, Dynamic instability of microtubules as an efficient way to search in space. *Proc. Natl. Acad. Sci.* **91**, 5682–5685 (1994).

3. C. Brossard, V. Feuillet, A. Schmitt, C. Randriamampita, M. Romao, G. Raposo, A. Trautmann, Multifocal structure of the T cell – dendritic cell synapse. *Eur. J. Immunol.* **35**, 1741–1753 (2005).

4. A.-K. Weier, M. Homrich, S. Ebbinghaus, P. Juda, E. Miková, R. Hauschild, L. Zhang, T. Quast, E. Mass, A. Schlitzer, W. Kolanus, S. Burgdorf, O. J. Gruß, M. Hons, S. Wieser, E. Kiermaier, Multiple centrosomes enhance migration and immune cell effector functions of mature dendritic cells. *J. Cell Biol.* **221**, e202107134 (2022).

5. F. Verde, M. Dogterom, E. Stelzer, E. Karsenti, S. Leibler, Control of microtubule dynamics and length by cyclin A- and cyclin B-dependent kinases in Xenopus egg extracts. *J. cell Biol.* **118**, 1097–1108 (1992).

6. R. Wollman, E. N. Cytrynbaum, J. T. Jones, T. Meyer, J. M. Scholey, A. Mogilner, Efficient Chromosome Capture Requires a Bias in the 'Search-and-Capture' Process during Mitotic-Spindle Assembly. *Curr. Biol.* **15**, 828–832 (2005).

7. G. Letort, F. Nedelec, L. Blanchoin, M. Théry, Centrosome centering and decentering by microtubule network rearrangement. *Mol. Biol. Cell* **27**, 2833–2843 (2016).

8. S. Som, S. Chatterjee, R. Paul, Mechanistic three-dimensional model to study centrosome positioning in the interphase cell. *Phys. Rev. E* **99**, 012409 (2019).
